# Molecular and Morphological Signatures of Chordate Development: Two Distinct Pathways, One Tunicate

**DOI:** 10.1101/801589

**Authors:** Mark Kowarsky, Chiara Anselmi, Kohji Hotta, Paolo Burighel, Giovanna Zaniolo, Federico Caicci, Benyamin Rosental, Norma F Neff, Katherine J Ishizuka, Karla J Palmeri, Jennifer Okamoto, Tal Gordon, Irving L Weissman, Stephen R Quake, Lucia Manni, Ayelet Voskoboynik

## Abstract

All chordates, including urochordates such as tunicates, develop through embryogenesis. The chordate larvae of colonial tunicates metamorphose to lose all chordate structures such as notochord, neural tube, segmented musculature, and then develop by asexual reproduction [blastogenesis], whereby stem cells form tissues and organs. These two developmental pathways establish the same body axis, morphogenetic patterning and organ formation. It is unknown if this convergent morphology implies convergent cellular and molecular mechanisms, and whether the stem cells that mediate these processes differ. Using the colonial tunicate *Botryllus schlosseri*, we combined transcriptome sequencing and multiple microscopy techniques to study the molecular and morphological signatures of cells at each developmental stage of embryogenesis and blastogenesis. This revealed that the molecular programs are distinct, but the blastogenic tissue-specific stem cells and embryonic precursor populations share similar molecular profiles. By comparing embryogenesis in other chordates we found shared developmental principles, highlighting transcription factors as key evolutionary conserved elements. This study establishes a platform for advancing the science of stem cell biology and regulation of development and regeneration.

## Introduction

One of the most challenging questions in developmental biology is how a complex multicellular organism develops from a single cell. During embryogenesis, spatio-temporal expression patterns progress while a single totipotent cell differentiates and divides into the various lineages that comprise the adult body (Conklin, 1905, Sulston et al., 1983). Studies across deuterostome species revealed numerous highly conserved patterning pathways and transcription factors (Davidson et al., 2002, Farley et al., 2015, Krumlauf 1994, Prummel et al., 2019, Simakov et al. 2015, Wodarz and Nusse 1998). Recently, advances in single-cell sequencing and lineage tracing have allowed high precision molecular and cellular timelines of the stem and progenitor cells involved in embryonic development and adult stem cell maintenance to be produced for chordates (Briggs et al. 2018, Cao et al., 2019, Chan et al., 2018, Farrell et al., 2018, Nowotschin et al., 2019, Schaum et al., 2018, Wagner et al., 2018). Other developmental pathways including asexual reproduction and whole body or tissue regeneration differ in origin but similarly establish body axes, morphogenetic patterning and organs (Fincher et al., 2018, Gerber et al., 2018, Manni and Burighel, 2006, Sabbadin et al., 1975, Voskoboynik et al., 2007). Although studies have identified evolutionarily conserved aspects of development (Duboule, 1994; Hu et al., 2017; Irie and Kuratani, 2011), embryogenesis has not been linked to asexual reproduction pathways like blastogenesis or other developmental modes. In particular, it is unknown if convergent morphology implies convergent molecular mechanisms, how organogenesis differs during sexual (embryogenesis), asexual and regenerative processes, and how the stem cells mediating these processes vary (i.e. tissue specific or pluripotent cells that differentiate).

We have found that colonial tunicates are the key to answering these questions. These marine organisms are unique among chordates: they can produce its adult body every week through two pathways (Figure S1), as well as possessing whole body regenerative capabilities (Gasparini et al., 2014, Manni et al., 2019, Voskoboynik et al., 2007). The adult stage of the colonial tunicate *Botryllus schlosseri* contains many individuals (zooids), derived by asexual reproduction from a single metamorphosed larva. These clonal progeny exist within a gelatinous tunic connected by an extracorporeal blood vessel system (Figure 1 and S1). Sexual development begins with fertilization of an egg by free-swimming sperm creating a zygote that develops over the next six days inside the zooid (at 18°C), ultimately releasing a swimming larva (Video1). That larva features chordate characteristics such as a notochord, neural tube and segmented musculature. Upon settling, the larva metamorphoses into an invertebrate individual, an oozooid (zooid originating from oocyte) which already has initiated asexual reproduction via buds, the precursors for the next generation’s zooid (blastozooid, zooid originating from buds). This oozooid begins a weekly asexual budding cycle termed blastogenesis in which secondary buds grow into primary buds which in turn complete organogenesis and replace their parent zooid (Figure S1; Video 2). At the end of the weekly cycle, the parent zooids in the colony undergo a synchronized programmed cell removal and are resorbed and cleared through phagocytosis, a process that eliminates most cells excluding the stem cells (Laird et al., 2005, Rinkevich et al., 2013, Rosental et al., 2018, Voskoboynik et al., 2008). Neighboring colonies can recognize each other and either form an immune rejection barrier between them or fuse blood vessels to form a single multilineage colony (Sabbadin, 1962, Scofield et al., 1982). Self/non-self recognition is governed by a single, highly polymorphic histocompatibility locus [called BHF] where colonies that share one or both BHF alleles are compatible (Voskoboynik et al., 2013a). After fusion, stem cells from each individual participate in the formation of new organs within developing buds, competing for germline and/or somatic lineages (Laird et al., 2005, Rinkevich et al., 2013, Rosental et al., 2018, Sabbadin and Zaniolo, 1979, Stoner et al., 1996 and 1999, Voskoboynik et al., 2008). Heritable germline stem cell predation potential leads to all gametes coming from one genome, even though the somatic tissues are natural chimeras (Sabbadin and Zaniolo, 1979, Stoner et al., 1996 and 1999, Weissman 2000 and 2015). The BHF histocompatibility polymorphism limits germline predation mainly to kin. These properties make *B. schlosseri* an ideal model for studying stem cell competition limited by histocompatibility, and mechanisms of regeneration (Manni et al., 2019, Voskoboynik et al., 2013b, Weissman 2000 and 2015).

**Figure 1:**
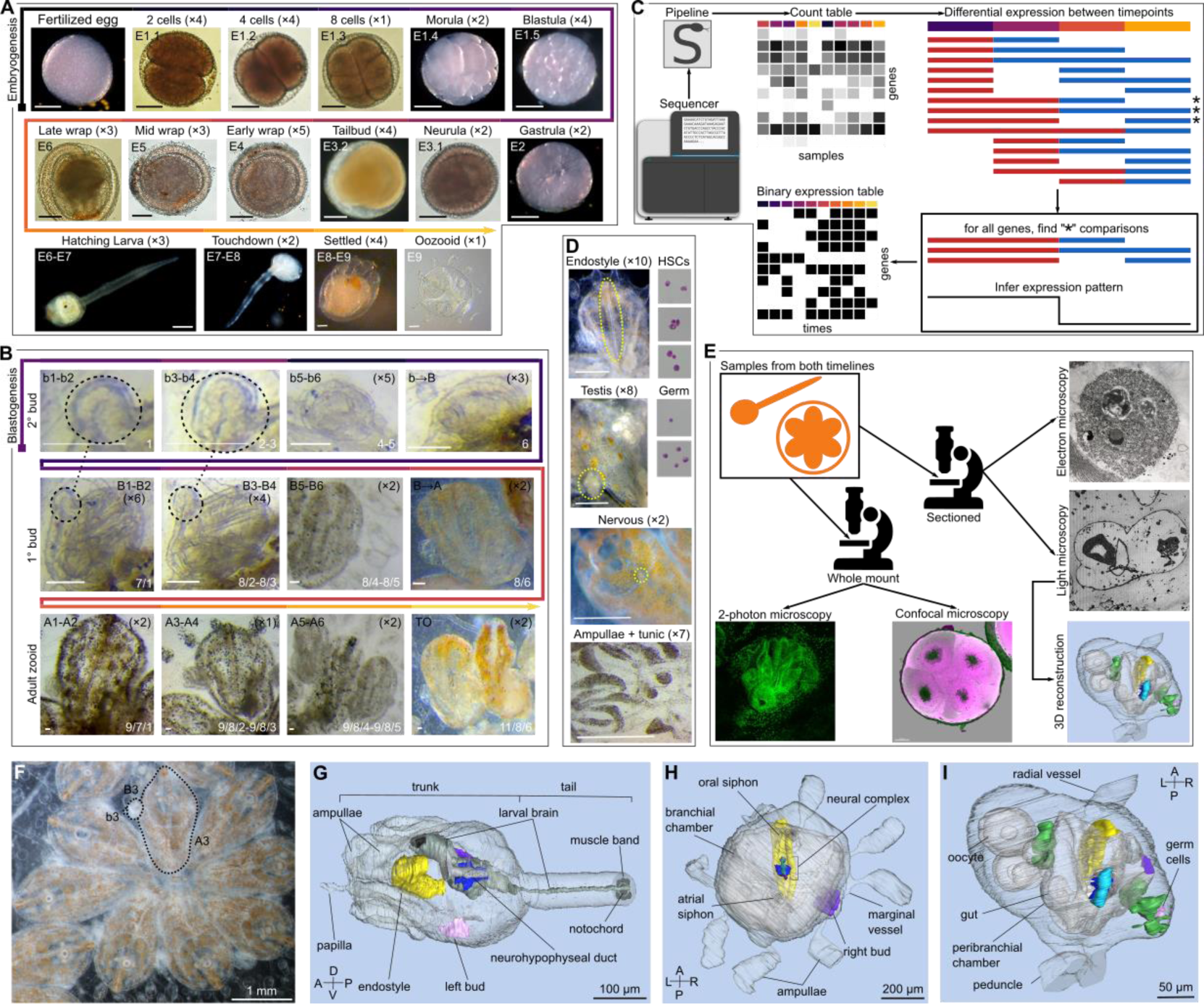
Sampling and methods for compiling the *B. schlosseri* atlas. A) The embryonic pathway from a fertilized egg until after metamorphosis. Numbers in parentheses after the staging name indicate the number of samples used in the transcriptomic analysis. Labels on the images indicate the numbering scheme used, where E indicates embryogenesis and the first number is the number of days typically elapsed during development at 18-20 °C. The second digit (where present) counts the stages present in a single day of development. Scale bars are 100 μm for the top two rows, and 60μm for the lower row. B) The blastogenic pathway for three generations (secondary buds (b), primary buds (B) and adult zooids (A)). Dotted circles and lines indicate that samples were collected together. Labels on images are similar to for embryogenesis, with numbers indicating the days of development of that generation, or transitions between generations. The white labels on the lower left of the images are the staging names according to Sabaddin (Sabbadin, 1955). Scale bars are 100μm. C) Description of pipeline for determining developmentally dynamic genes. Reads from the sequencer are processed using a Snakemake pipeline to produce a gene count table for each gene. Samples from different sets of time (red vs blue) are compared for differential expression. For each gene, all the comparisons that were significant (*) are gathered, and the optimal binary expression pattern is inferred, producing a binary expression table for the different timepoints. D) Various tissues and cell types used in the transcriptome analysis. E) Building a morphological atlas for sexual and asexual development including 3D reconstructions from histological serial sections. F) *B. schlosseri* colony with dotted lines showing an adult (A3), with a primary bud (B3) and secondary bud (b3) attached to it. G-I) 3D reconstruction of (G) a larva in early metamorphosis (E8) from transverse histological serial sections; (H) an oozooid (E9) from oblique histological serial sections, with its secondary bud at advanced stage (b4); (I) a bud (B1) with its secondary bud (b1) from oblique histological serial sections. The bud is joined to the colonial circulatory system through a radial vessel and to the parent through a stalk like part (peduncle). Tunic is omitted in all the 3D reconstructions; the outer epithelium is the epidermis. Labelled larval structures are: cerebral ganglion (blue), dorsal organ (burgundy), dorsal tube (light blue), endostyle (yellow), germ cells (light pink), larval nervous system (dark grey), left bud (stage b1, light violet), neural gland (green), neural gland aperture (purple), neurohypophyseal duct (light gray), photolith (black), right bud (advanced stage b1, dark violet), testis (green). Other structures are transparent. Dorsal view.

To compare the molecular and morphological signatures associated with embryogenesis and blastogenesis, we studied developing embryos and buds at key sequential developmental stages (Figure 1A-B). We generated comprehensive whole transcriptome sequence data and developed a bioinformatic method to identify developmentally dynamic genes via their expression (Figure 1C). Coupling this with confocal, two photon, light and electron microscopy (Figure 1E), we characterized the stages of embryonic and blastogenetic development and built an atlas of sexual and asexual development [*Tabula compositi chordati*] (Tables S1-S4). This study outlines the molecular and morphological landscape of these two developmental modes to provide candidates for an integrative understanding of the pathways that direct embryogenesis and blastogenesis. The comparison revealed that multiple molecular paths can lead to the same outcome (Figure 2). Using tissue and cell-type specific molecular signatures (Figure 1D and S2; Table S3) with detailed morphological data, that includes three-dimensional reconstructions, we identified and compared the developmental origin during embryogenesis and blastogenesis of the central nervous system (Figure 3 and S3), the hematopoietic system (Figure 4 and S4), the endostyle (Figure 4 and S4) and the reproductive systems (Figure 5 and S5). We found equivalent tissue specific transcriptional timings in both developmental pathways, excluding the nervous system (Figure 6 and S6). Our findings suggest that cellular developmental trajectory is defined early, and demonstrates that the blastogenic tissue specific stem cells and embryonic precursor populations share similar molecular dynamics and organogenesis timing (Figures 3-6 and S3-S6). Comparing *B. schlosseri* developmental pathways to amphioxus and zebrafish embryogenic transcriptomes (Farrell et al., 2018, Marlétaz et al., 2018), revealed that many transcription factors are evolutionarily conserved elements of chordate development (Figure 7 and S7). The experimental approach and computational tools we developed provide an efficient and powerful way to produce and analyze complex molecular and morphological data. The atlas and resources crafted for this project can be employed to study regulation of development and regeneration processes in complicated systems including mammals.

**Figure 2:**
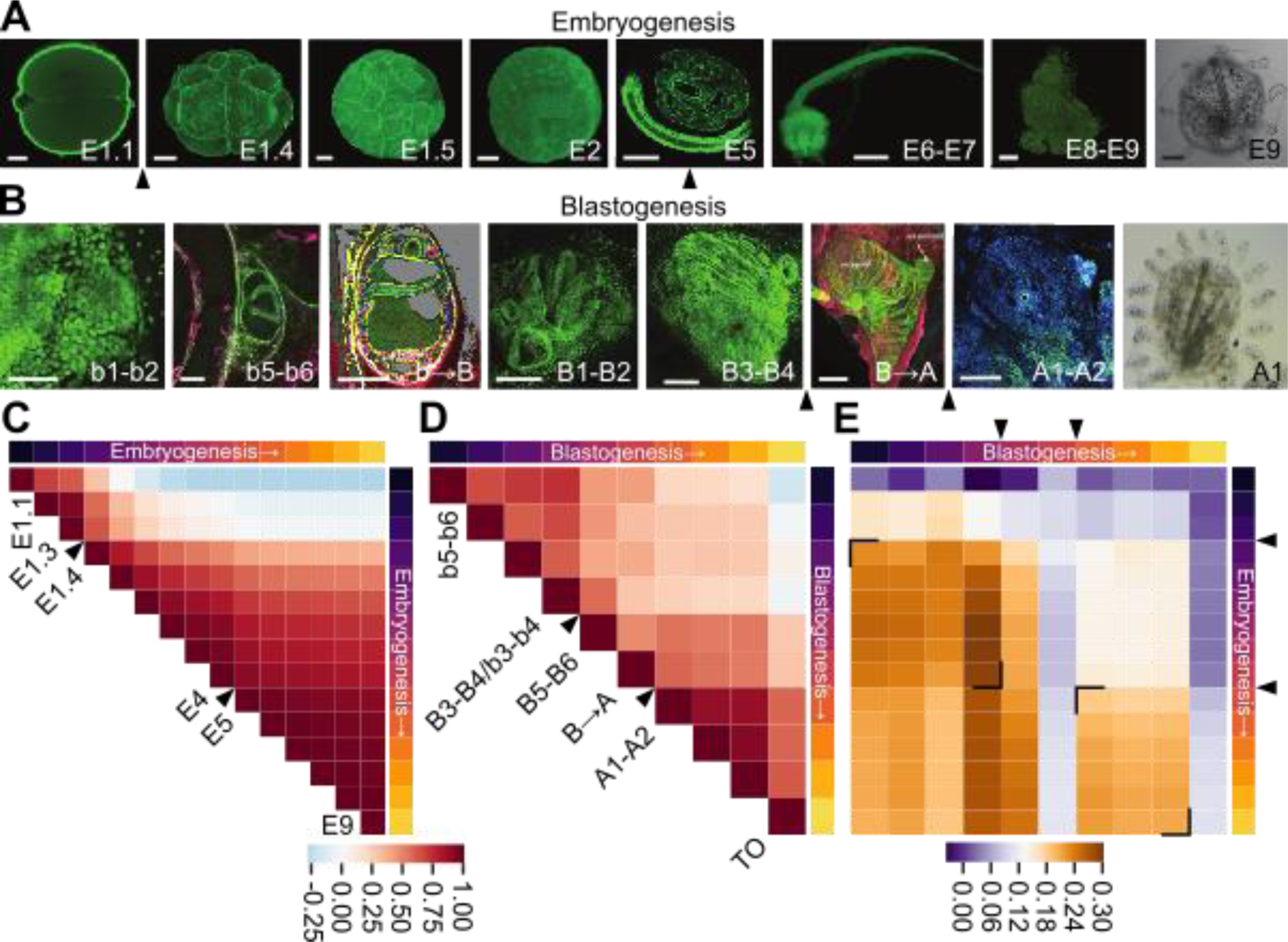
Morphological and molecular transitions in embryogenesis and blastogenesis. A) Images of select embryonic stages: two cells (E1.1), morula (E1.4), blastula (E1.5), gastrula (E2), early wrap (E5), hatching larva (E6-E7), a settled larva undergoing metamorphosis (E8-E9) and an oozooid (E9). All except the last are confocal images, which is a live image. Scale bars in μm: 30, 30, 30, 50, 100, 60, 50, 70. B) Images of select blastogenic stages: the secondary bud (b1-b2, b5-b6, b→B), primary bud (B1-B2, B3-B4, B→A) and adult zooid (A1-A2) and live image of the blastozooid. Two photon images: b1-b2, B1-B2, B3-B4, A1-A2; confocal images: b5-b6, b→B, B→A. Scale bars in μm: 40, 50, 40, 100, 50, 500, 500. Colors: blue-nucleus, green-actin, red-cytoplasm and tunic. C) Correlation of genes enriched at different times in embryogenesis. Black arrowheads indicate transitional times between transcriptionally similar eras. D) Same as (C), but for blastogenesis. E) Correlation between time signatures across the two developmental pathways. Black corners indicate regions of similarity.

**Figure 3:**
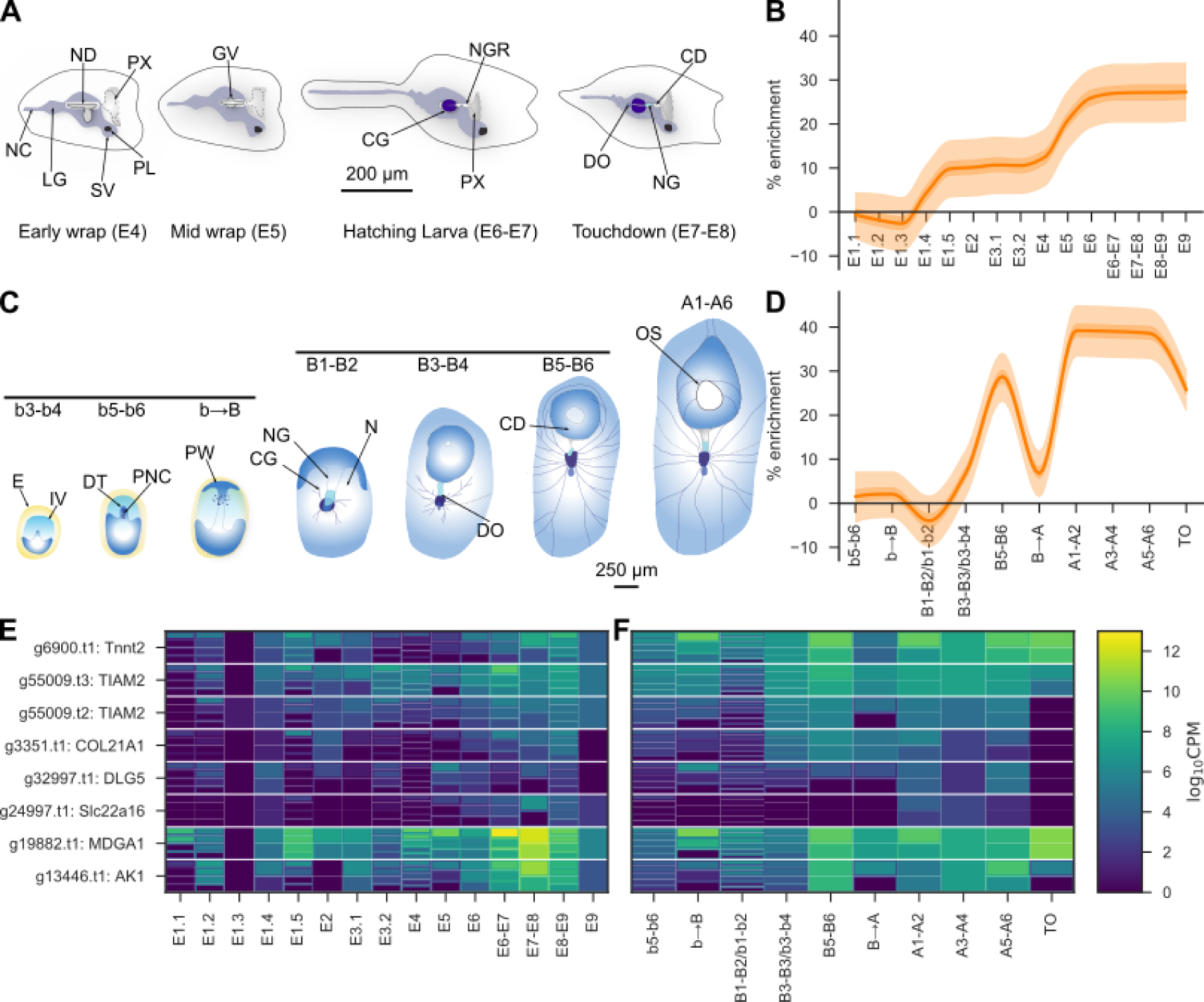
Origin and development of the nervous system. A) Schematic illustration of nervous system development during embryogenesis (early wrap (E4) to settled larva (E7-E8)). Labels: NC-nerve cord, LG-larval ganglion, SV-sensory vesicle, PL-photolith, PX-pharynx, ND-neurohypophyseal duct, GV-ganglionic vesicle, NGR-neural gland rudiment, CG-cerebral ganglion, CD-ciliated duct, DO-dorsal organ, NG-neural gland. B) Gene enrichment plot of central nervous system-associated genes during embryogenesis. Baseline (0%) is set under the null model of a random subset of genes. Light and dark shaded regions indicate the 50% and 99% confidence intervals under a hypergeometric model. C). Schematic illustration of nervous system development during blastogenesis (secondary bud (b3-b4) to an adult zooid (A1-A6). Labels: E-epidermis, IV-inner vesicle, DT-dorsal tube, PNC-pioneer nerve cells, PW-pharyngeal wall, CG-cerebral ganglion, NT-neural gland, N-nerves, DO-dorsal organ, CD-ciliated duct, OS-oral siphon. D) Gene enrichment plot of central nervous system-associated genes during blastogenesis. E-F) Klee plots of transcription factors found to be dynamic between E4-E7 that are expressed in the nervous system and associated in the literature with nervous system development/differentiation. Klee plots are heatmaps that allow for heterogeneous groupings of samples (i.e. different numbers of samples at a given time point). Around and between each grouping the mean value of the group is shown in the outline. E) embryogenesis. F) blastogenesis.

**Figure 4:**
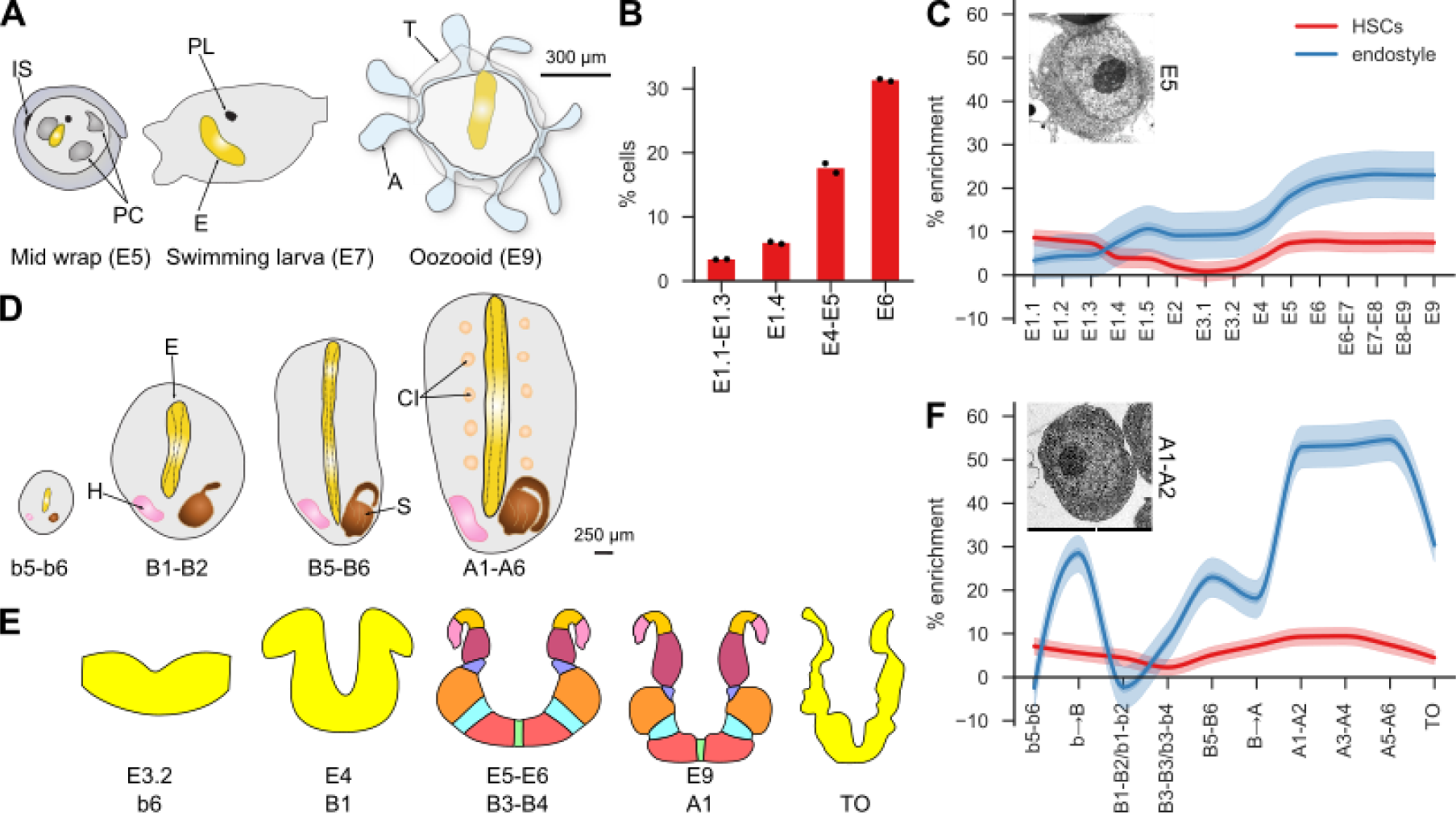
Origin and development of hematopoiesis and its niche (endostyle). A) Schematic illustration of the endostyle development during embryogenesis (mid wrap (E5) to oozooid (E9)). Labels: IS-interpapillary space, PC-peribranchial chamber, PL-photolith, E-endostyle, T-tunic, A-ampulla. B) Proportion of cells in embryos that are identified as enriched for HSCs using a FACS based assay. C) Gene enrichment plot of HSC and endostyle associated genes during embryogenesis. Inset is TEM image of a hemoblast after they become visible. D) Schematic illustration of endostyle development during blastogenesis (b5-b6 - A1-A6). Labels: H-heart, E-endostyle, S-stomach, CI-cell islands. E) Illustration of the transverse section of the endostyle during development in both embryogenesis and blastogenesis showing the formation of the eight zones. Details in Figure S4. F) Gene enrichment plot of cells enriched for HSC and endostyle associated genes during blastogenesis. Inset is a hemoblast visible in an adult zooid showing similar morphology to that in embryogenesis.

**Figure 5:**
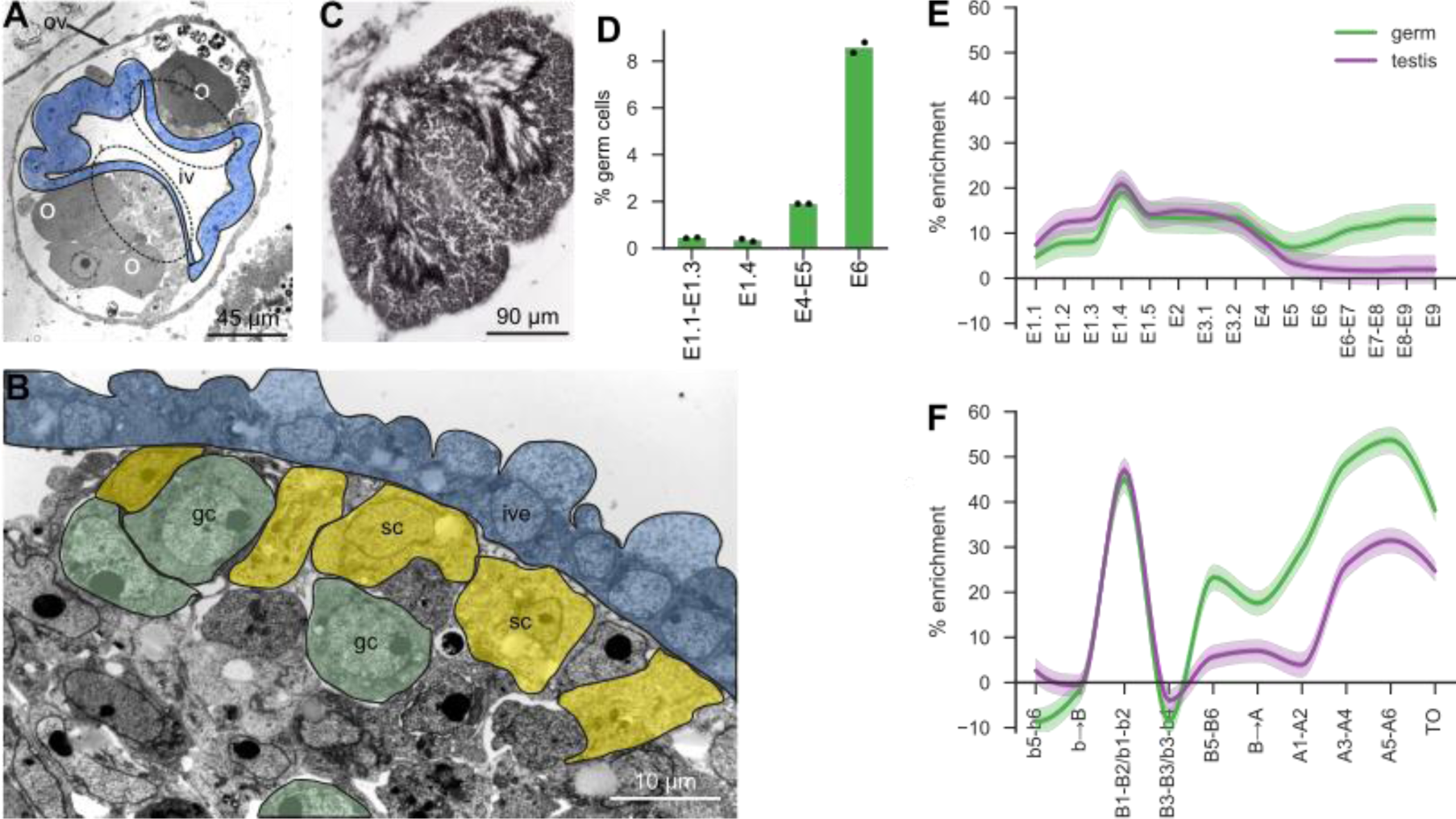
Origin and development of germ cells and the testis. A) Transverse section of secondary bud (b5) with bilateral gonad rudiment (dotted lines) pressing the inner vesicle (iv, blue) and detail of the gonadal blastema. Some previtellogenic oocyte (O) are close to the epidermis, the outer vesicle (ov). Toluidine blue. B) Candidate male primordial germ cells (gc, green) are grouped together with presumptive somatic cells (sc, yellow). They are close to the inner vesicle epithelium (ive, blue), TEM. C) Section of two testis lobules belonging to an adult zooid. Immature germ cells are at testis periphery, whereas mature sperms are central. Toluidine blue. D) Proportion of cells in embryos that are identified as germ cells using a FACS based assay. E-F) Gene enrichment plot of germ and testis associated genes during embryogenesis (E) and blastogenesis (F).

**Figure 6:**
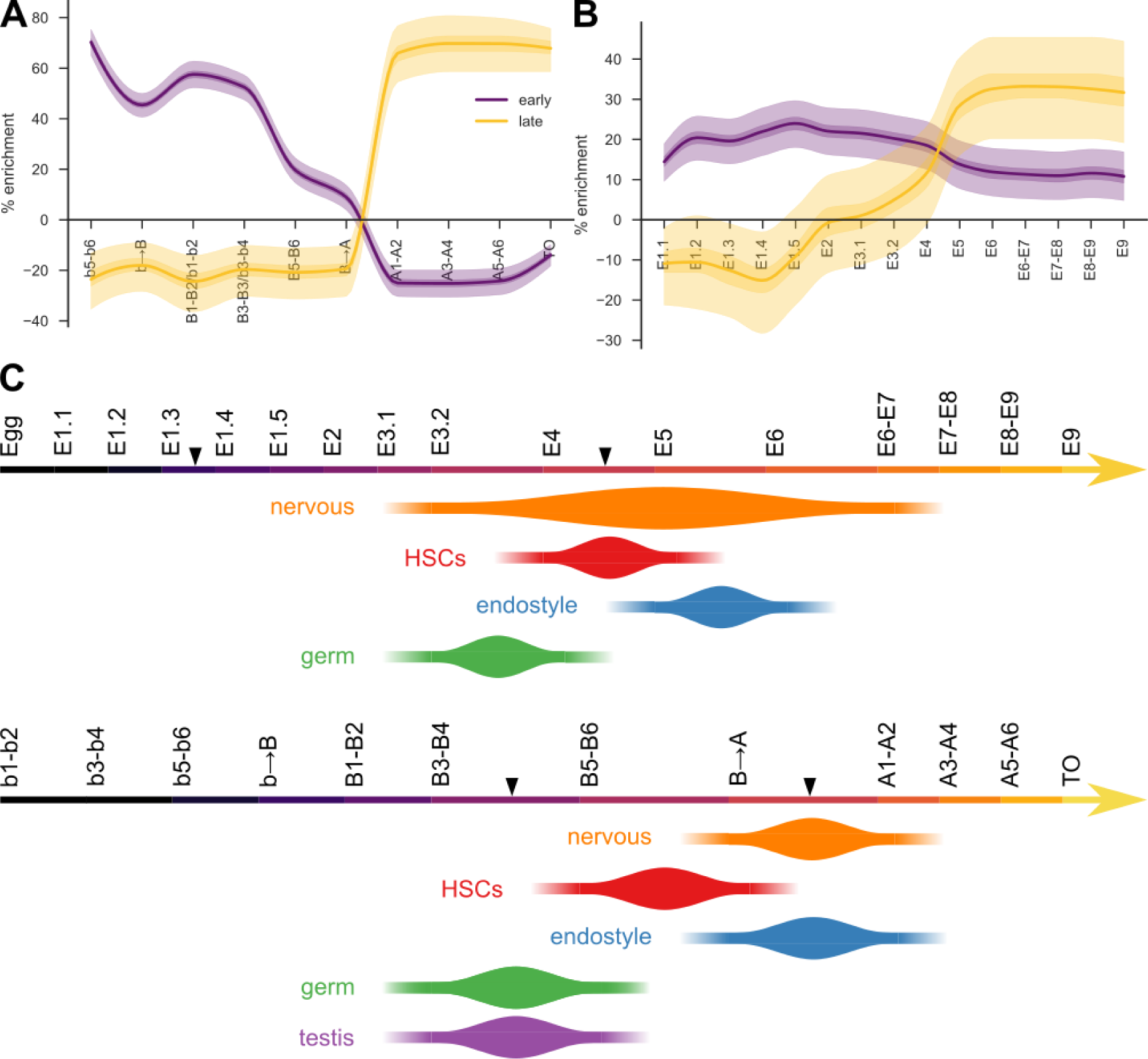
Timeline of organogenesis inferred from transcriptional signatures. A) Gene enrichment plot of early (expressed only before B→A) and late (expressed only after B→A) HSC-associated genes during blastogenesis. Baseline (0%) is set under the null model of a random subset of genes. Light and dark shaded regions indicate the 50% and 99% confidence intervals under a hypergeometric model. B) Gene enrichment plot of the same sets of genes as shown in panel (A), but during embryogenesis. C) Estimated regions in organ development of the transition between early/late shared gene expression. Arrowheads on axes indicate the transitions between eras found based on whole transcriptome profiling (figure 2C-D).

**Figure 7:**
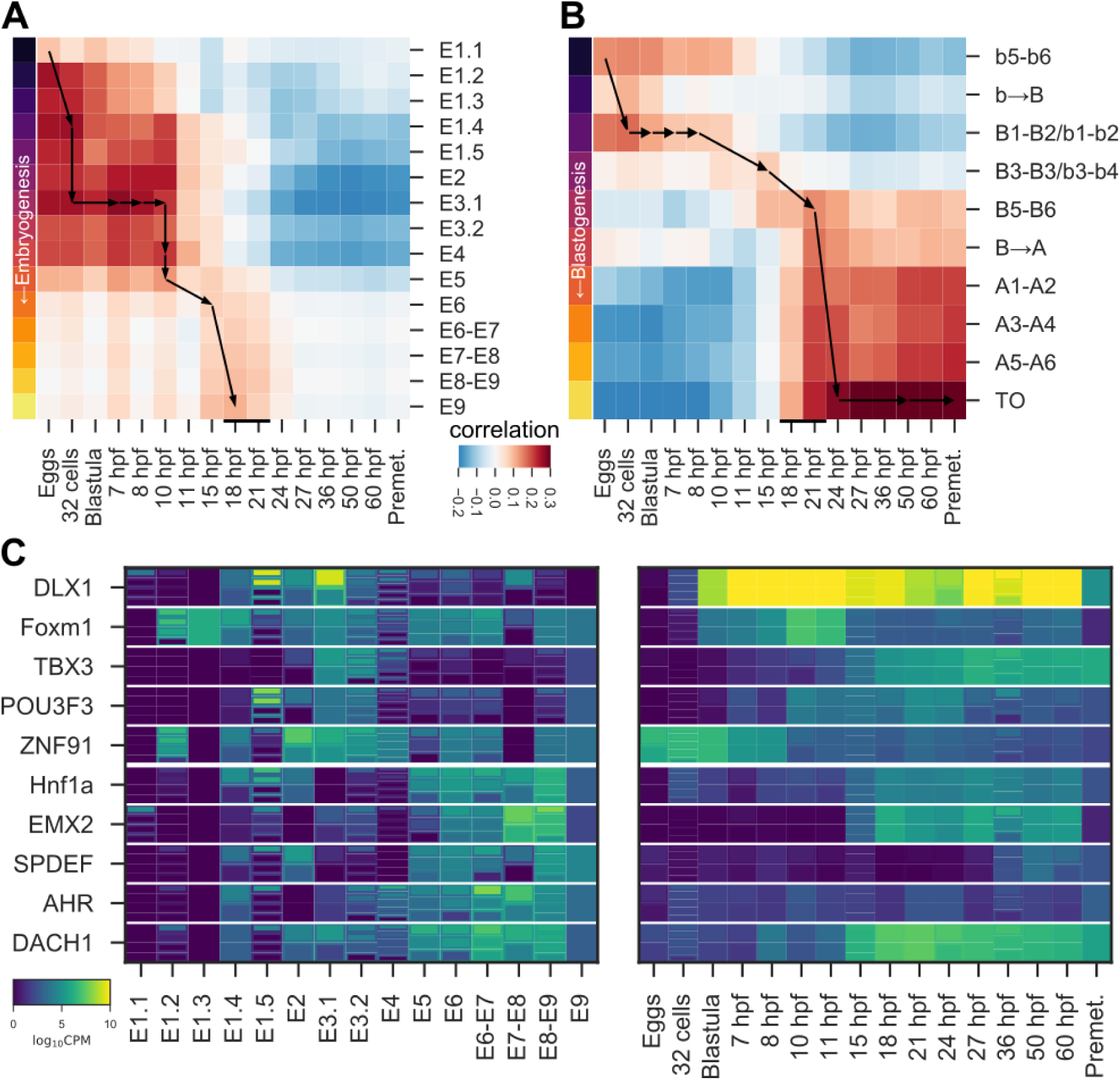
Comparison between *B. schlosseri* developmental pathways and amphioxus embryogenesis. A) Correlation between shared transcription factors at various time points between embryogenesis in *B. schlosseri* (rows) and in amphioxus (columns). Arrows indicate the estimated path of equivalent times in the two pathways. The phylotypic period in amphioxus is 18-21 hpf and indicated by a thin black bar. B) Like (A), but for blastogenesis in *B. schlosseri*. C) Klee plot of the top (by fold change) transcription factors shared between species (*B. schlosseri* and amphioxus) that are co-expressed either early (upper 5) or late (lower 5) (see Figure 2E) in embryogenesis and blastogenesis in *B. schlosseri*.

## Results

### Building a Molecular and Morphological Atlas of Two Developmental Pathways

To characterize the molecular and morphological pathways that define *B. schlosseri* embryogenesis and blastogenesis, samples were collected at different developmental stages. For embryogenesis, specimens from 15 different developmental stages across a time frame of 7 days, were taken for RNA extraction and sequencing (Figure 1A; n=44). For blastogenesis, we sequenced secondary buds, primary buds, combinations of primary and secondary buds and the whole adult zooid from 12 different developmental stages across a period of 7 days (Figure 1B; n=29). Corresponding samples were taken for bright-field, histology, confocal, two photon, and electron microscopy to build the morphological atlas of sexual and asexual development (Figure 1E). To identify tissue/cell specific signatures, we sequenced different tissues and organs from adult individuals including: testes, endostyles, brains (cerebral ganglion and neural gland), circulating enriched stem cell populations (hematopoietic; candidate germline stem cell), and blood vessels embedded within the tunic (Figure 1D). We generated comprehensive whole transcriptome sequence data from a total of 73 samples (multiple biological replicates, see annotations in Figure 1A-B, D) taken during the selected key sequential developmental stages.

A detailed morphological description of all stages was compiled (Table S1) based on confocal, 2 photon microscopy, serially cut histological sections and electron microscopy (EM) (transverse, sagittal, and frontal plane; 1 μm). For both developmental pathways a numerical staging method that refers to the day of development for embryo, secondary bud, primary bud and adult zooid was used (Table S1 and Figure S1), simplifying the comparison between *B. schlosseri*’s two developmental pathways and embryogenesis in other chordate species. A detailed description of organ development (Table S3) through embryogenesis and blastogenesis was obtained by EM (Figures 3-5 and S3-S5). We made three dimensional reconstructions of a larva in early metamorphosis (E7-E8; Figure 1G), an oozooid (E9; Figure 1H), and a primary bud and secondary bud (B1+b1; Figure 1I) using 426, 853, and 375 sections respectively. These reconstructions revealed 19 anatomical entities in the larva, 17 in the oozooid, and 31 in the buds, which can be viewed using the morphodynamic browser MorphoNet (Leggio et al., 2019).

Following sequencing, reads were processed using a Snakemake (Köster and Rahmann, 2012) pipeline (see methods) resulting in a transcript count table (Table S4). We identified developmentally dynamic genes by finding all differentially expressed genes between sets of contiguous times in a pathway using edgeR (Robinson et al., 2010) (Figure 1C, upper right). For each gene, times with statistically significant differences (FDR < 0.05) were recorded. By comparing different time signatures the pattern that best supported the observed differential expressions for each gene was selected (Figure 1C, lower right). To further simplify the comparisons, these time signatures were binarized, with 1 indicating “high” expression and 0 indicating “low” or zero expression producing a gene-time expression matrix for each gene along the developmental pathways. The developmentally dynamic genes are presented in Table S2A-B.

### Embryogenesis and Blastogenesis have Distinct Molecular and Morphological Programs

Embryogenesis and blastogenesis have vastly different morphological patterns that nonetheless produce similar adult bodies (Figure 2A-B and S1, Table S1). Here we compare how the overall molecular programs for these two processes are related, to determine where the molecular pathways converge and where they differ. For each developmental program, we calculated the correlation of the binary developmentally dynamic gene matrix at each stage against all other stages to determine periods of similar and distinct gene expression in an unbiased manner. This analysis revealed several “eras” of development separated by transcriptional transition stages (Figure 2C and 2D, Table S2C).

In embryogenesis, the highest degree of similarity was between neighboring times as would be expected by the overall gradual and unidirectional nature of this process (Figure 2C). This analysis revealed three major eras of embryogenesis: (i) two to eight cells (E1.1-E1.3), (ii) morula to early wrap (E1.4-E4), and (iii) mid wrap to the oozooid (E5-E9). Similarly in blastogenesis, the greatest transcriptional correlations occurred between neighboring times (Figure 2D), but in contrast to embryogenesis it has a discontinuous trend. This trend was due to both our inability to sample the early stages of blastogenesis (secondary buds b1-b4) without parental tissue, and the longer time periods between sample collection compared to embryogenesis (Figure 1B). This analysis revealed two main eras and several transitional stages of blastogenesis: (i) from the fifth developmental day of the secondary bud to the fourth developmental day of the primary bud and secondary bud (b5-b6 to B3-B4/b3-b4), and (ii) the adult zooid from the time the siphons open (A1) through to takeover (TO). The stages at the end of the first era (B5-B6 to B→A) did not cluster with other stages instead creating a distinct transitional time (Figure 2D). The genes specific to these time periods that have human or mouse gene homologs, were analyzed using Gene Analytics (Ben-Ari Fuchs et al., 2016). This allowed us to investigate the top pathways and gene ontologies associated with each time period in a systematic manner.

The first embryonic era, characterized by the first three cleavages, is typical in all tunicates (Conklin, 1905, Hotta et al., 2007, Stach and Anselmi, 2015), and expresses a unique molecular profile that significantly differs from later stages (Figure 2C). We hypothesize that this is in part due to the maternal-to-zygotic transition (Lee et al., 2013), the period in which the developing embryo begins to generate its own transcripts rather than those inherited maternally from the egg. Amongst the top pathways expressed in this era are genes associated with metabolism of proteins and gene expression. Approximately 400 transcription factors (TFs) are activated during this period, including pioneer zinc finger and nuclear receptor TFs key to development and regulation, as well as TFs associated with stem cell pluripotency and circadian rhythm like *Prox1*, *GSC* and *ATF4*.

The second era begins at the morula stage leading into the formation of three germ layers, and the development of the chordate neural plate and neural tube closure. The era concludes with the separation of the tail and trunk regions (tailbud) where the following organs and tissues are present: heart, digestive system, branchial and peribranchial chambers, larval brain, notochord, nerve cord, rudiment of the adult neural complex, striated muscles and ampullae (Figure 2A, Table S1A and S3). Many TFs that were not expressed during the first era are activated in the second era, including approximately 400 TFs associated with: gene expression, chromatin organization, mesodermal commitment pathway, neural crest differentiation, regulation of pluripotency, and Wnt/Hedgehog/Notch pathway. The top expressed pathways are chromatin regulation, mRNA splicing, gene expression and metabolism.

The third era begins at the mid-wrap stage and ends when the larva completes its development and metamorphoses into an oozooid (Figure 1G-H and 2A). The larva has oral and atrial siphons, a branchial sac with stigmata and endostyle; a digestive system with an esophagus, stomach and intestine; a larval brain containing a photolith as well as the adult neural complex; three sensory papillae, eight ampullae, two buds, and a tail. All organs remain after metamorphosis excluding the tail, anterior papillae and the larval brain (Figure 1H and Table S1A). Fifty new TFs are activated during this era, including those associated with cell differentiation, neuron fate specification, heart development, thyroid gland development, epithelial cell differentiation, and organ morphogenesis. All previous pathways that were highly expressed in the first and second eras excluding metabolism, are down regulated. Pathways associated with homeostasis and maintenance are highly expressed in this era. There are low numbers of genes associated with differentiation and high numbers associated with remodeling and large structural changes in the developing organism. During metamorphosis low numbers of molecular changes are detected, in agreement with morphological data that shows minor organ loss in the transition from larva to oozooid (Figure 1G-H).

During the first era of blastogenesis all the organ rudiments form within the secondary bud (Figure 1I and Table S1). The secondary buds are derived from the peribranchial epidermis which closes to form a vesicle (inner vesicle), surrounded by the parental epidermis (outer vesicle) (Manni et al., 2014). Circulating haemocytes originating from the colony home to the buds (Rinkevich et al., 2013, Rosental et al., 2018, Voskoboynik et al., 2008). During this era all the organ rudiments are formed including: branchial and peribranchial chambers, intestine, heart, nervous system, and in fertile colonies the gonads. The first era concludes when the heart begins to beat in the primary bud. The top pathways involved in this era are: gene expression, mRNA splicing, cell cycle and mitosis. Approximately 600 TFs associated with gene expression, chromatin regulation, hematopoietic stem cell gene regulation, mesodermal commitment, and mitochondrial gene expression are expressed, reflecting the massive development occurring during this era.

During the transitional period between eras in the primary buds (B5-B6 to B→A), there continues to be a high expression of genes associated with cell cycle and mitosis along with the expression of pathways associated with organelle biogenesis and lipid metabolism. During this period the expression of about 200 TFs that were expressed in the first era are down regulated while TFs associated with gene expression, cell differentiation, nervous system development, adipogenesis, Wnt/Hedgehog/Notch, pluripotency, TGF beta signaling, and neural crest remain active. This is consistent with the morphological data that shows the primary bud continuing to grow in dimension while the bud’s organs complete their differentiation and specification. These final differentiation steps prepare the primary bud for its transformation into a functional filter feeding adult zooid.

In the second era, the adult zooid undergoes relatively minor growth and morphological changes. The main events characterizing this era are ovulation, sperm release and zooid regression. The replacement of a zooids’ generation occurs at the end of this era (takeover=TO), with all zooids in a colony degenerating in a synchronized wave of massive apoptosis and phagocytic cell removal (Ballarin et al., 2008, Lauzon et al., 1993). A similar transcriptional profile is expressed by zooids during stages A1 to A6 while the TO profile differs from the zooid prior to take over (Figure 2D). During this era 370 TFs are expressed, but only 108 TFs are expressed during TO (Table S2B) which include five zinc finger TFs, *HNF1a*, *RFX6* and *ZKSCAN1*. Other highly expressed pathways during the second era include those associated with integrin, adhesion, lipid metabolism, pathways associated with cardiac hypertrophy and those associated with degradation of the extracellular matrix.

The top features characteristic of the developmental eras in embryogenesis and blastogenesis are highly dissimilar to each other despite the shared body plan and same organs and tissues that develop. To directly compare them, we calculated the correlation of activated developmentally dynamic genes in the same manner as was done above within a single developmental pathway (Figure 2E). While these two pathways both express similar numbers of genes, they are highly dissimilar, having a maximum correlation of only 0.3. Interestingly, genes that are co-expressed are less likely to be *B. schlosseri* specific, indicating that shared developmental genes are more likely to be conserved between species than those in a specific pathway. There are 389 TFs unique to embryogenesis, 76 unique to blastogenesis and 576 that are shared.

The first embryonic era (E1.1-E1.3) is distinct from all blastogenic stages. This is expected, considering that this era contains a process exclusive of initial embryogenesis cleavage. Likewise, the periods during takeover (B→A and TO) have transcriptional profiles distinct from that of any embryonic stages (Table S2C). Pathways associated with toll-like receptors, innate immune response and inflammation, as well as pathways associated with gene expression are highly expressed during these time points (Table S2C). This is in part expected, since they are events marking a discontinuity in an individuals’ life that does not have any correspondence in embryogenesis. Surprisingly there is no correspondence with metamorphosis, during which larval tissue resorption is also governed by apoptosis (Karaiskou et al., 2015). Although the two pathways can be considered transcriptionally distinct, some of the genes highly expressed during embryogenesis are also highly expressed during blastogenesis (Figure 2E and Table S2C).

There are two blocks of stages with shared expression: (i) embryonic stages of morula (E1.4) to early wrap (E4) match the blastogenic stages of secondary bud (b4-b5) to primary bud and secondary bud (B3-B4/b3-b4), and (ii) embryonic stages from mid wrap (E5) to oozooid (E9) match the adult zooids (A1-A6) excluding the takeover stage. From a morphological point of view (Table S1A), the first block of stages shows the organization of the body plan, whereas the second block regards stages where the adult organs have comparable mature states. In each of these blocks, the genes that were most strongly co-expressed in both pathways were selected and investigated for functional enrichment (Table S2C). Early in development, pathways linked to chromatin regulation, gene expression, mRNA splicing, telomere extension and the mitotic cell cycle are dominant. Late in development, pathways associated with metabolism, degradation of the extracellular matrix, collagen chain trimerization and transport of glucose dominate. This supports a model that cells are dividing and differentiating into organs early in development, whereas late in development mostly homeostasis is taking place. We hypothesize that stem cell activity would be higher earlier on, and indeed, when we looked at the expression of genes known to regulate pluripotent stem cells we found 13 that were co-expressed in the early time periods vs only three that were co-expressed in the later time periods. The early expressed genes include homologs to *Wnt5a*, *Fzd2*, *NR6A1* and *Smad2*.

### Organ Enriched Molecular Profiles Correlate with Morphogenesis

To identify and compare the developmental origin of specific organs and tissues during embryogenesis and blastogenesis we studied the nervous system, the hematopoietic system, the endostyle and the reproductive system, using histological data and tissue and cell-type specific transcriptome libraries (Figure S2, Table S2D and S3). For each system and organ, the relative enrichment of genes associated with that tissue was compared along each developmental pathway. We compared regions of change to concurrent morphological changes to begin to identify and analyze the genes that may drive these changes and identify transcription factors that may precede or drive organ development.

#### Origin of the nervous system

The *B. schlosseri* larva has a part of the central nervous system specific for it, and in addition form the rest of the nervous system that degenerates and regenerates each blastogenic cycle (Table S3). In the adult, the brain consists of an ovoid cerebral ganglion, located dorsally between the oral and atrial siphons, possessing a cortex of neuronal somata, an inner medulla of neurites (Figure 1H and S1K-L) and several mixed nerves emanating into the organism (Burighel and Cloney, 1997, Manni and Pennati, 2015). The cerebral ganglion resides by the neural gland, a sac-like structure open anteriorly into the pharynx and together they constitute the neural complex (Manni and Pennati, 2015). In the embryo, the oozooid brain develops concurrently with the larval brain. The larval brain develops from the typical chordate neural plate, sharing several homologies with the vertebrate brain (Cao et al., 2019) (Figure 3A). The larval brain is active during the motile stage, when the larva hatch, find an open subtidal surface, co-settle with siblings with a bias for cosettlement of histocompatible offspring to allow chimera formation, indicating olfactory-like sensory input (Grosberg and Quinn, 1986). The larval brain is connected to a photolith which has 2 photoreceptor cells, indicating other sensory inputs limited to the chordate stage. The adult brain is retained after metamorphosis in the oozooid, functions during the sessile stage, where it controls siphons, branchial sac’s stigmata, and the body muscle’s contractions (Figure 3C). It may also control *B. schlosseri’*s synchronized blastogenic cycles where the zooids degenerate on cue, while in the primary and secondary buds neurogenesis creates *de novo* a new adult neural complex. It would be surprising if any type of long term memory function could be inherited by the bud, inasmuch as it is regenerated from stem/progenitor cells anew each cycle.

The origin of the nervous system in mammals begins with the induction of the neural ectoderm shortly before gastrulation (Spemann and Mangold, 2001) and was tracked to the 32 cell stage (morula) in the solitary tunicate, *Ciona intestinalis* (Nishida and Stach, 2014). The first enrichment of nervous system genes in *B. schlosseri* occurs between 8 cells (E1.3) and blastula (E1.5) (Figure 3B). The enrichment profile stabilizes during the gastrula, neurula and early tailbud stages, when the neural plate forms, and develops the main components of the larva nervous system (Figure 3A and S3A-H, Table S3). The second enrichment of genes appears during the wrap stages (E4-E6), when the development of the larval and oozooid brain become evident (Figure 3A and S3 A-H, Table S3). Out of 28 TFs that are activated on E1.3, nine are associated with neural crest, brain and nervous system development including Pou3f3, Dix1, Lmx1a, Sox6, PA2G4, Zmat4, Zfp318 Cebpz and Aedbp1. The expression of these TFs most likely induce the neural ectoderm in the *B. schlosseri* embryos at E1.3. There are 5273 genes that are dynamically expressed during the wrap stages (E4-E7), 78 are associated with the *B.schlosseri* nervous system and 27 of these have mammalian homologs. Of these, eight genes are known to be involved in neuron differentiation or nervous system development including *TIAM2*, *DLG5* and *MDGA1* that are involved in regulating cell proliferation, neurite growth, migration and axon guidance. The expression of these genes are shown in the Klee plots (Figure 3E-F) along both developmental pathways. Their expression increased during key nervous system developmental stages during both embryogenesis and blastogenesis. During metamorphosis and the following stages, the gene signature again stabilizes. During these stages the larval nervous system is resorbed while the cerebral ganglion remains (Figure S3E-H).

The larval nervous system is composed of a large sensory vesicle with a photolith, a small ganglionic vesicle, a visceral ganglion, a neck, and nerve cord (Manni et al., 1999). The photoloith can detect light and gravity receiving input from two photoreceptor cells, a statocyst and other specialized cells (Sorrentino et al., 2000). In solitary tunicates on the other hand, the larvae photoreceptor cells are located in the ocellus and the statocyst in the otolith (Ryan et al. 2016). Many genes known to be expressed in the retina and photoreceptor pigmented epithelium of mammalian eyes are highly expressed in the larva of *B. schlosseri* (Table S2A). After metamorphosis, the photolith is observed in the oozooid, however, it is resorbed and does not develop in the asexual generation of buds that will replace the founder oozooid. The oozooid cerebral ganglion forms from a tubular structure, the neurohypophyseal duct, that derives from the anterior neural plate (Table S3). The duct, which represents the rudiment of the entire neural complex, is initially connected to the ganglionic vesicle but later on it loses this original connection, and grows forward to open into the anterior pharynx. The pharynx aperture will form the ciliated duct of the neural gland, whereas its posterior part will form the body of the neural gland and the dorsal organ, considered homologous to the dorsal strand of other tunicates (Manni and Pennati, 2015). Meanwhile, the neurohypophyseal duct wall proliferates pioneer nerve cells, which coalesce to organize the cerebral ganglion.

During blastogenesis, the neural complex rudiment forms from a thickening of the dorsal area of the secondary bud (b5-b6) (Figure S3I-J). This thickened area develops into the dorsal tube, which grows forward and fuses with the anterior pharyngeal wall during the transition into the primary bud (b→B). At the same time, pioneer nerve cells appear on the dorsal tube while the dorsal tube loses its posterior connection with the forming atrial chamber and differentiates into the neural gland. In the early primary bud (B1-B2) the cerebral ganglion and the neural gland are connected, pioneer nerve cells proliferate and motor nerves appear (Figure S3K-L). Then (B3-B4) the neural gland separates from the posterior dorsal organ and numerous nerves appear (Figure 3M). When the primary bud is about to replace the resorbing zooid (B→A), the dorsal organ is entirely isolated from the neural gland, and a massive reduction in the number of nerves occurs. In the adult zooid, the cerebral ganglion is organized into cortex and medulla (Figure S3N); these structures are progressively destroyed during TO (Figure S3O-P), when apoptosis leads to processes such as phagocytosis [efferocytosis of dying cells; programmed cell removal of live cells] reabsorbs the brain, together with all other tissues.

A high degree of concordance between known differentiation timelines, morphological observations and tissue-associated transcriptional profiles are observed in blastogenesis (Figure 3D). In particular, the proportion of nervous system-associated genes defined as active at each blastogenic stage reflects the nervous system’s development and morphogenesis processes. Such as the reduction of nerves observed in the primary bud during generation changes (B→A) (Figure 3D) which is accompanied with inhibition of 48 TFs associated with mammalian nervous system and brain development including SOX9, OTX2, FOXD3, and SMAD1 (Table S2B).

Taken together, *B. schlosseri* offers an unique system to study neurogenesis: During chordate embryonic development a larval stage specific brain is formed with functions required for a swimming larva to complete its passage from ‘hatching’ to settlement on a subtidal surface, after which the larval brain is resorbed during the extensive asexual reproductive cycles of blastogenesis. The embryonic phase also developed a second brain, connected with the larval brain, but which persists after metamorphosis: this brain is regenerated during every blastogenic cycle, perhaps analogous with mammalian neurogenesis from persisting central nervous system stem cells.

#### Origin of the Hematopoietic System and its Niche

In the mammalian blood system, all mature blood lineages, including erythrocytes, platelets, and all innate and adaptive immune cells, are generated from hematopoietic stem cells (HSCs) (Spangrude et al., 1988). Hematopoiesis and the factors that regulate HSC self-renewal are highly conserved between vertebrates (Jagannathan-Bogdan and Zon, 2013). Recently we identified HSCs and progenitor cells in *B. schlosseri* (Rosental et al., 2018). Adult blood cells in *B. schlosseri* include cells with morphologies similar to lymphocytes (including HSCs, myeloid lineage cells and perhaps other immune cells), phagocytic cells (hyaline amoebocytes and macrophage-like cells), morula cells (large granular lymphocyte like precursors and activated morula cells) pigmented cells and nephrocytes (Ballarin and Cima, 2005, Rosental et al., 2018). In embryogenesis blood cells with similar morphological features are found (Burighel et al., 1983, Milanesi and Burighel, 1978). EM analysis of embryos revealed a gradual emergence of morphologically similar blood cells (Table S3). First, lymphocyte-like cells with a high nucleus-cytoplasm ratio, rich with ribosomes as well as morula cells are detected in early wrap (E4) (Figure S4A-B and Table S3). By mid-late wrap (E6), hyaline amoebocytes with small membrane-bound granules in the cytoplasm, and pigment cells are detected (Figure S4C-E). During metamorphosis, macrophage-like cells with vacuoles containing heterogeneous content appear (E7-E8) (Figure S4F) and nephrocytes containing granules with hourglass geometry are detected in the oozooid (E9) (Figure S4G).

The enrichment of *B. schlosseri* HSC associated genes during embryogenesis revealed an increase in their expression from E1.1 to E1.4 (Figure 4C), including 239 genes with human homologues expressed in hematopoietic bone marrow and 43 with human homologues expressed in HSCs. In the solitary tunicate *Ciona*, haemocytes are recognized only after metamorphosis. In Ciona haemocytes originate from the trunk lateral cells (visible in the tailbud period), whose lineage has been traced to the 64-cell embryos (cleavage period) (Kawaminani and Nishida, 1997), corresponding to E1.4-E1.5 in *B. schlosseri*. The *Botryllus* HSC gene enrichment signature is high early in development (E1.1-E1.4), decreases during E1.5-E3, increases again during the wrap stages (E4-E6), then stabilizes after the larval stage (Figure 4C). The expression of a subset of these genes, including *ALOX5*, *SELE* and *SELP* homologs have expressions about 1000-fold higher in late wrap than early wrap. Morphologically, the latter increase coincides with the appearance of hemoblasts (for a morphological term that includes candidate HSCs) in the early wrap (E4) (inset in Figure 4C). This cellular characterization was validated by flow cytometry (Figure 4B).

The HSC enrichment pattern shows a relatively steady signature along blastogenesis, with lower expression in primary and secondary buds on developmental days 3 and 4 (B3b3-B4b4) and in zooids during TO (Figure 4D). While the molecular signature of hemocytes circulating in the relatively small secondary bud during b5-b6 was detected, this signature is diluted in the primary and secondary buds in stages B3b3-B4b4. As the number of HSCs and their progenitors increase in the primary buds and zooids during B5 to A5, the enrichment of genes selectively expressed in these HSCs increases as well. Towards the TO stage the HSC signature decreases (Figure 4B).

During blastogenesis stem cells including HSCs apparently reside and proliferate in the anterior ventral side of the endostyle (Figure S4K-O) (Rosental et al., 2018, Voskoboynik et al., 2008). The endostyle is a long glandular groove extending medially at the ventral face of the zooid branchial sac along its anterior posterior axis consisting of eight distinct symmetric anatomical zones (Figure S4N). It is immersed in blood flow through the large subendostylar sinus and other sinuses (Burighel and Brunetti, 1971) (Figure S4Q-R). In embryogenesis, the endostyle is first recognized during mid wrap (E5, Figure 4A-E and S4H-J), coincident with the increase of associated genes (E4-E6), which succeeded those of HSCs (E4) (Figure 4B). However, in blastogenesis, unlike HSCs the endostyle gene enrichment fluctuates dramatically (Figure 4F). The first wave of gene enrichment (b5-b6 in Figure 4F) is associated with the appearance of the endostyle (Table S3, Figure 4D,E, and S4K). The formation of the 8 zones (Figure 4D-E and S4M-N) coincides with the second wave of gene enrichment (B5-B6, A1) (Figure 4F). During takeover when the endostyle is destroyed (Figure S4O), the gene enrichment drops off (Figure 4F).

In both embryogenesis and blastogenesis, the endostyle follows the same developmental stages creating a highly similar anatomy with eight zones (Figure 4E). Extending medially along the anterior posterior axes of both the primary buds and early wrap embryos and expressing TF essential to early development including FOXD3, PPARG, ATF4, PROX1 PPARD (Table S2C).

#### Origin of Reproductive Tissues

The germ line is made up of a highly protected and strictly regulated group of cells that transmit genetic information to the next generation. It was initially proposed (Weismann, 1892), and later shown that they originate as a very small founding population that is segregated from somatic cells early in development (Anderson et al., 1999, Dixon, 1994, Soriano and Jaenisch, 1986; Ueno et al., 2009).

There is a large degree of correlation between the enrichment profile of germ and testis associated genes not surprisingly, as almost 30% of the enriched genes are shared (Figure 5E and F, Table S2D). In embryogenesis (Figure 5E), the two transcriptional signatures track each other well until the late wrap stage (E6). The oozooid is not sexually mature, so while general stem-associated genes continue to be expressed (germ signal), those for the testis decrease to the baseline. There is a slow increase in the germ stem cell signal during early cleavage (E1-E3); at these stages, candidate germ cells expressing vasa were identified in *B. schlosseri* (Brown et al., 2009). The initial peak in the morula (E1.4) aligns with high expression of *NR6A1* (aka *GCNF*-germ cell nuclear factor), a transcription factor involved in germ cell development and neurogenesis and also corresponds to the emergence of germline stem cells at the 4- to 16-cell stage in mice (Anderson et al., 1999, Dixon, 1994, Soriano and Jaenisch, 1986).The peak also precedes, when germ cells are first recognized in other tunicates: the 64-cell stage, in *Halocynthia roretzi* (Kawamura et al., 2011), and the gastrula (stage 13) in *Ciona* (Shirae-Kurabayashi et al., 2006). The increase enrichment of the germ gene signature after E6 is reflected in flow cytometry measurements (Figure 5D).

*B. schlosseri* is a hermaphrodite and gonads are detected in mature colonies only after several blastogenic generations (Kawamura et al., 2011). Each adult individual usually possesses male and female organs on both the lateral body wall.

In blastogenesis (Figure 5F), the male gonad rudiment appears in the secondary bud (b3) in the form of a loose mass of primordial germ cells and somatic cells close to the inner vesicle wall (Table S3). Candidate male germ cells with high nucleus cytoplasm ratio, one or more nucleoli, several ribosomes, and mitochondria are observed during this stage (Figure 5A-B and S5A). Somatic cells, which later are involved in testis wall formation, are close to germ cells, often at the periphery of the germ mass. Germ cells expressing vasa were detected at this stage (Rosner et al., 2009). As soon as the testis wall is formed (b5), the testis becomes a compact mass in which spermatogonia undergo meiosis. This wall separates the testis from the forming female gonad (when present) and subdivide it into lobes (testicular follicles), in which an initial lumen can be seen (Figure S5B-C). In the primary bud (Figure S5D-E), the testis progressively enlarges and completes its development. The mature testis in adult zooids possesses 4-5 lobes displaying a gradient of sperm maturation: spermatogonia are located at lobe periphery, followed by primary spermatocytes, secondary spermatocytes, spermatids, and fully mature spermatozoa at the lobe center (Figure S5F-H); sperm are released at stage A3.

Germ and testis enrichment profiles both show sharp and strong enrichment in the primary buds (b1-b2/B1-B2) (Figure 5F). This corresponds to the time immediately before gonad establishment in secondary buds (b3), and the emergence of testis gross anatomy in primary buds (B3). A second sharp increase in testis gene expression enrichment characterizes the stages of late primary bud, involved in spermatid maturation, and the adult zooids at stage A3-A4, corresponding to the last phases of testis maturation preceding spawning. Germ cell enrichment is very high in the adult stages, they constitute a pool of cells able to circulate in the colony vasculature, reside in cell islands (Figure S4Q-T), and colonize the gonad niches in the following blastogenic generations (Rinkevich et al., 2013, Sabbadin and Zaniolo, 1979). In chimeras between BHF compatible colonies, the germline cells are almost exclusively derived from one of the parent colonies in the chimera, and this is heritable both by asexual and sexual reproduction (Stoner and Weissman, 1996, Stoner et al, 1999].

Many markers for primordial germ cells are observed during early embryogenesis and blastogenesis including: alkaline phosphatase (Chiquoine 1954, Ginsburg et al., 1990), SSEA-1/FUT4 (Marani et al., 1986), Oct3/4/POU (Rosner et al., 1990), Blimp-1/PRDM (Saitou et al., 2005), Piwi (Cox et al. 2000) and vasa/DDX4 (Raz 2000; Castrillon et al. 2000). These shared signatures along these two developmental pathways forms a link between the germline embryonic and adult stem cells.

#### Tissue Specific Transcriptional Timings are Shared in Both Developmental Pathways

An open question is to what extent organ development in each pathway shares molecular signatures. In order to answer this, for the developmentally dynamic tissue-associated genes, we systematically defined those that were enriched early and late in each pathway and compared the enrichment of those genes in the other pathway (Figure S6). This resulted in identifying both the time that early → late gene patterns transitioned as well as the correspondence of organogenesis times between the two pathways (Figure 6 and S6, Table S3). For example in blastogenesis there is a clear early/late pattern of HSC associated gene enrichment that is switched during the B → A stage (Figure 6A). The same pattern for this gene list is also observed in embryogenesis at E4-E5 (Figure 6B). When applied to all tissue/cell types examined in this study, this analysis revealed both the conservation of early/late tissue specific gene expression (molecular development) and the shared chronology of tissue emergence (global development) (Figure 6C).

These expression patterns can help find the genes associated with stem versus differentiated cell state during organogenesis. Our findings suggest that cellular trajectory is defined early in development and demonstrates that blastogenic tissue specific stem cells and their embryonic precursor cells share similar molecular dynamics (e.g. hematopoietic and germline stem cells, germline stem cells).

### Evolutionary Conserved Transcription Factors Regulate Chordate Development

To place *B. schlosseri* development in the context of other chordates we compared its developmentally dynamic genes with those in amphioxus (*Branchiostoma lanceolatum*) and zebrafish (*Danio rerio*) using publicly available data (Marlétaz et al., 2018) (Figure 7 and S7). For these other species, the developmentally dynamic genes were calculated in the same way as for *B. schlosseri*. Comparing the embryogenesis timelines between the amphioxus and zebrafish using our method confirms the location of the phylotypic period, the period with maximal molecular similarity (Duboule, 1994; Hu et al., 2017; Irie and Kuratani, 2011), as similar to that found in the literature (Marlétaz et al., 2018).

Comparing all pathways (*Botryllus* embryogenesis, *Botryllus* blastogenesis; amphioxus embryogenesis, zebrafish embryogenesis), we focused on either evolutionarily conserved putative transcription factors (Figure S7A) or all genes with the same annotations (Figure S7B). The estimated equivalence of developmental trajectories tends to have a more consistent and continuous downward trend only when the enriched transcription factors are correlated (Figure S7A). By comparison, when all genes are compared (Figure S7B), the developmental trajectories tend to be less consistent (for example, amphioxus/zebrafish vs *B. schlosseri* embryogenesis). This supports the idea that expression and timing of transcription factors is conserved in development, but that the genes activated/repressed downstream are less evolutionarily constrained. Early *B. schlosseri* embryogenesis (E1.2-E4) resembles that of early embryogenesis in other chordates: amphioxus, egg-10 hours post fertilization (hpf) and zebrafish, 2-12 hpf (Figure 7A andS7). The final product of embryogenesis (oozooid, E9), does not align with latest embryogenesis stages of amphioxus and zebrafish. Instead it correlates with 18hpf in amphioxus (neurula stage), and 12-16 hpf in zebrafish (Figure 7A and S7). Blastogenesis on the other hand extends across more time points, with the later periods (A1 to TO) having the strongest resemblance to the latter stages of amphioxus development (Figure 7B). The patterns for zebrafish are similar (Figure S7).

While *B. schlosseri* embryogenesis and blastogenesis pathways shared some TFs and pathways with embryogenesis of other chordates, they are highly dissimilar, having a maximum correlation of 0.3 (Figure 7A-B; Table S5). Focusing on periods with conserved TFs expression revealed that during early chordate development TFs that are associated with regulation of cell cycle (FoxM1), stem cell pluripotency (TBX3) and axial patterning (DLX1) are high, whereas the TFs that are expressed during late stages of development are mainly associated with metabolism (HNF1A), adipogenesis (HAR) and cell fate determination (DACH1) (Table S5; Figure 7C)).

In conclusion, in an evo-devo perspective, we show that transcription factors are evolutionarily conserved elements of chordate development and that *B. schlosseri*, with its synchronized life cycle, is an important model to study regulation of development [oogenesis] and regeneration [blastogenesis] processes in complicated systems including mammals.

## Acknowledgments

We thank C. Lowe, T. Raveh, C. Patton, R. Voskoboynik, A. Olson, Y. Voskoboynik, P. Bump, J. Thompson, J. Lee, B. Compton, N. Myers, T. Naik for technical advice and help, and Dr. E. Faure for hosting our dataset on MorphoNet.org. This study was supported by NIH grants R56AI089968, R01AG037968 and RO1GM100315 (to I.L.W., S.R.Q., and A.V.), the Chan Zuckerberg investigator program (to I.L.W and A.V), the Virginia and D. K. Ludwig Fund for Cancer Research, a grant from the Siebel Stem Cell Institute and a Stinehart-Reed grant (to I.L.W.). L.M. was supported by PRIN - Prot. 2015NSFHXF. C.A was supported by a Postdoctoral Fellowship of the Larry L Hillblom foundation, by Stanford School of Medicine Dean’s Postdoctoral Fellowship, by Aldo Gini foundation Fellowship and Iniziative di Cooperazione Universitaria 2017 Fellowship of the University of Padova. B.R. was supported by a Postdoctoral Fellowship of the Human Frontier Science Program Organization LT000591/2014-L and NIH Hematology training grant T32 HL120824-03, NIH shared equipment grant, 1S10OD025091-01.

## Author Contributions

Conception and design: A.V., M.K., L.M., C.A.; mariculture and sample collection: K.J.I., K.J.P.; RNA isolation and library preparation: K.J.P., A.V.; sequencing: J.O., N.F.N.; sequencing analysis and development of analytical tools: M.K., S.R.Q.; light and electron microscopy: L.M., P.B., G.Z., F.C., C.A.; 3D reconstruction: L.M., F.C.; confocal microscopy: K.H., C.A.; two photon microscopy: T.G., A.V.; flow cytometry and sorting: B.R.; writing of manuscript: M.K., A.V., L.M., C.A., K.J.P., K.J.I., I.L.W.; technical support and conceptual advice: K.H., N.F.N., S.R.Q., I.L.W.

## Data Availability

Sequencing data can be found on the NCBI Sequence Read Archive under accession

*Tabula compositi chordati* which includes detailed morphological and molecular characterization of *B. schlosseri* sexual and asexual development can be found on google shared drive Supplementary Tables and Videos and on morphonet browser using the following links:

Table S1 https://drive.google.com/file/d/1yv7RX0Ssp_axkgzSF-N3C77aFARQytL0/view?usp=sharing

Table S2 https://drive.google.com/file/d/1_KZBmhDc0ORhJwDH5DDm8b66Cjh7uHfC/view?usp=sharing

Table S3 https://drive.google.com/file/d/1X4JDuksmbcijBC7Jt-5mKM2QceLOpq7R/view?usp=sharing

Table S4

Supplementary video 1 https://drive.google.com/open?id=1ULn9xOdep0BTbRui3MM1assikVz3KNZ0

Supplementary video 2 https://drive.google.com/open?id=1UhEYRhIIyMg-kcVDXvEWssfKvcpGgTwf

Three dimensional reconstructions

http://morphonet.org Login: Botryllus3D Password: oozooid

## Supplementary Data

**Figure S1:**
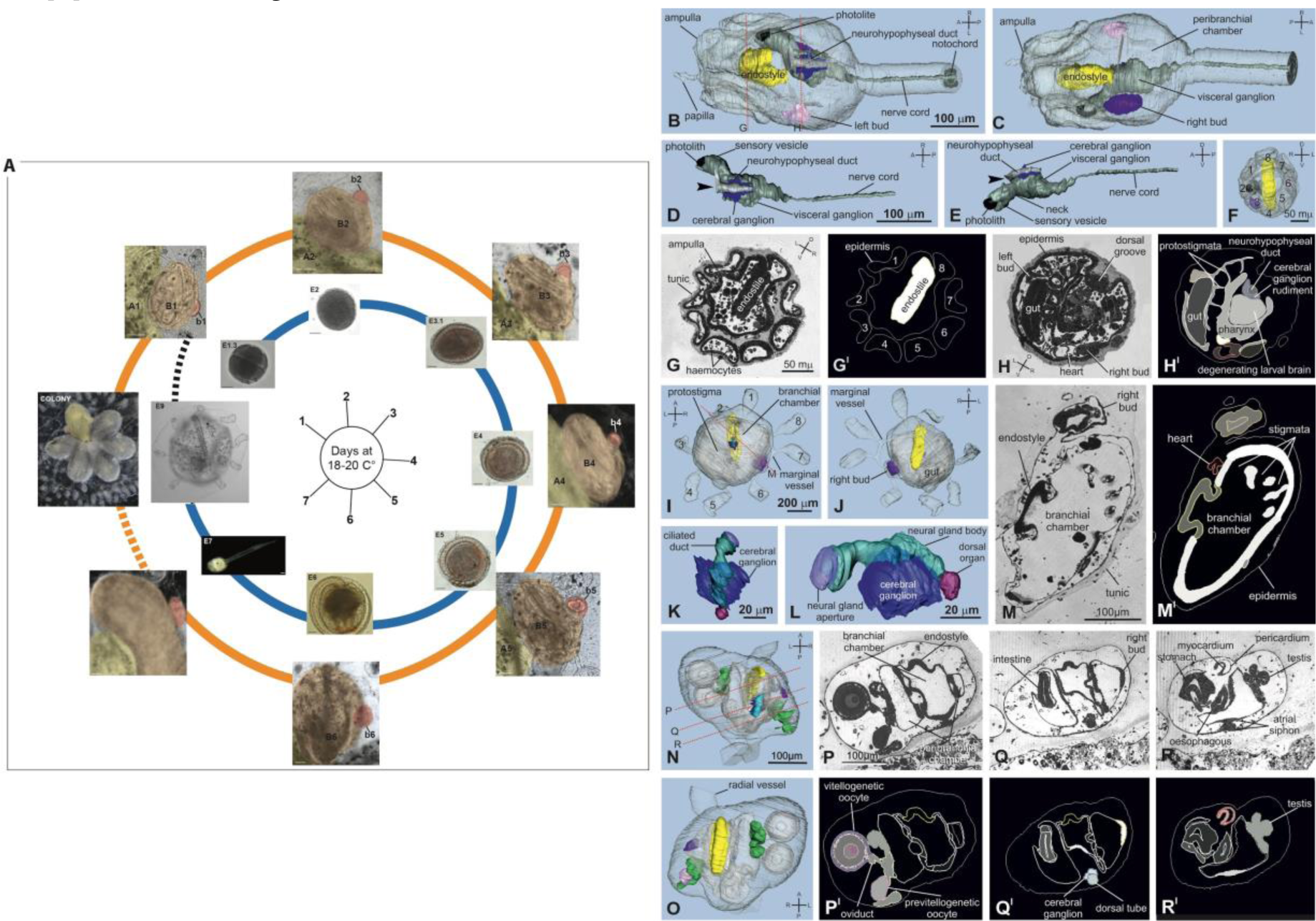
*Botryllus schlosseri* synchronized developmental pathways, and the anatomy of a larva, an oozooid, and a bud (3D reconstructions). A. Embryogenesis begins with the fertilization of an egg by sperm creating a zygote that develops over the next 6 days (18-20°C) inside the zooid, ultimately releasing a swimming larva (E7; day 7). The larva swims away from its mother colony, settles on a substrate (E8-E9), and metamorphoses into an oozooid (E9) which evaginates a primary bud which has an incipient secondary bud as the precursors for the next generation zooid. The weekly asexual budding cycle: begins with secondary buds (b1-b6) that grow into primary buds (b→B) which in turn complete organogenesis (B1-B6) and replace their parent zooid (B→A). Zooids open their siphons and live for six days (A1-A6), on day 7 all the zooids in the colony undergo a synchronized programmed cell death and removal and are resorbed and cleared through phagocytosis (takeover, TO). When a zooid grows more than one bud, a colony of genetically identical zooids is produced. (B-H) 3D reconstruction of a larva in early metamorphosis (E8) from transverse histological serial sections. Labeled larval structures are: budlets (the right one in dark violet, the left one in light violet), cerebral ganglion (blue), endostyle (yellow), larval nervous system (dark gray), neurohypophyseal duct (light gray), and photolith (black). Other structures are transparent; the tunic is not showed. The larva is viewed from the dorsal (B) and ventral (C) side. The red dotted lines in B represent the levels of sections in G and H. In D-E, dorsal (D) and right (E) views of the larval nervous system and the neural complex rudiment (neurohypophyseal duct and cerebral ganglion). F. Anterior view of the larva. Note the eight ampullae (1-8) for larval adhesion. G-H. Histological sections of the larva at the level of ampullae (G) and pharynx rudiment (H). Toluidine blue. G^I^ and H^I^ show the larval structures segmented on sections G and H. Note in H’ that the larval brain degenerates during metamorphosis; it is a structure that mediates perception and efferent signaling for the tadpole, not the blastozooid. (I-M) 3D reconstruction of an oozooid (E9) from oblique histological serial sections. Labeled structures are color coded as in B; further colored structures are: dorsal organ (burgundy), neural gland (green), neural gland aperture (purple). The eight ampullae (1-8) attach the oozooid to substrate. I: dorsal view; J: ventral view. The red dotted line in I represents the level of section in M. K-L. Neural complex viewed from dorsal (K) and lateral (L) side. M-M^I^.Histological section (M) and the corresponding segmented structures (M^I^). N-R) 3D reconstruction of a bud from oblique histological serial sections. Labeled larval structures are: budlets (the right one in dark violet, the left one in light violet), cerebral ganglion (blue), endostyle (yellow), larval nervous system (dark gray), neurohypophyseal duct (light bud at stage B1 with its right secondary bud at stage b1. Labeled structures are color coded as in B and I; further colored structures are: dorsal tube (light blue), germ cells (light pink) and testis (green). The bud is viewed from dorsal (N) and ventral (O) side. The red dotted lines in N represent the levels of sections in P-R. P-R. Histological sections of the bud at level of gonad (P), secondary bud (Q), and atrial siphon rudiment (R). Toluidine blue. P^I^-R^I^show the bud structures segmented on sections P-R.

**Figure S2:**
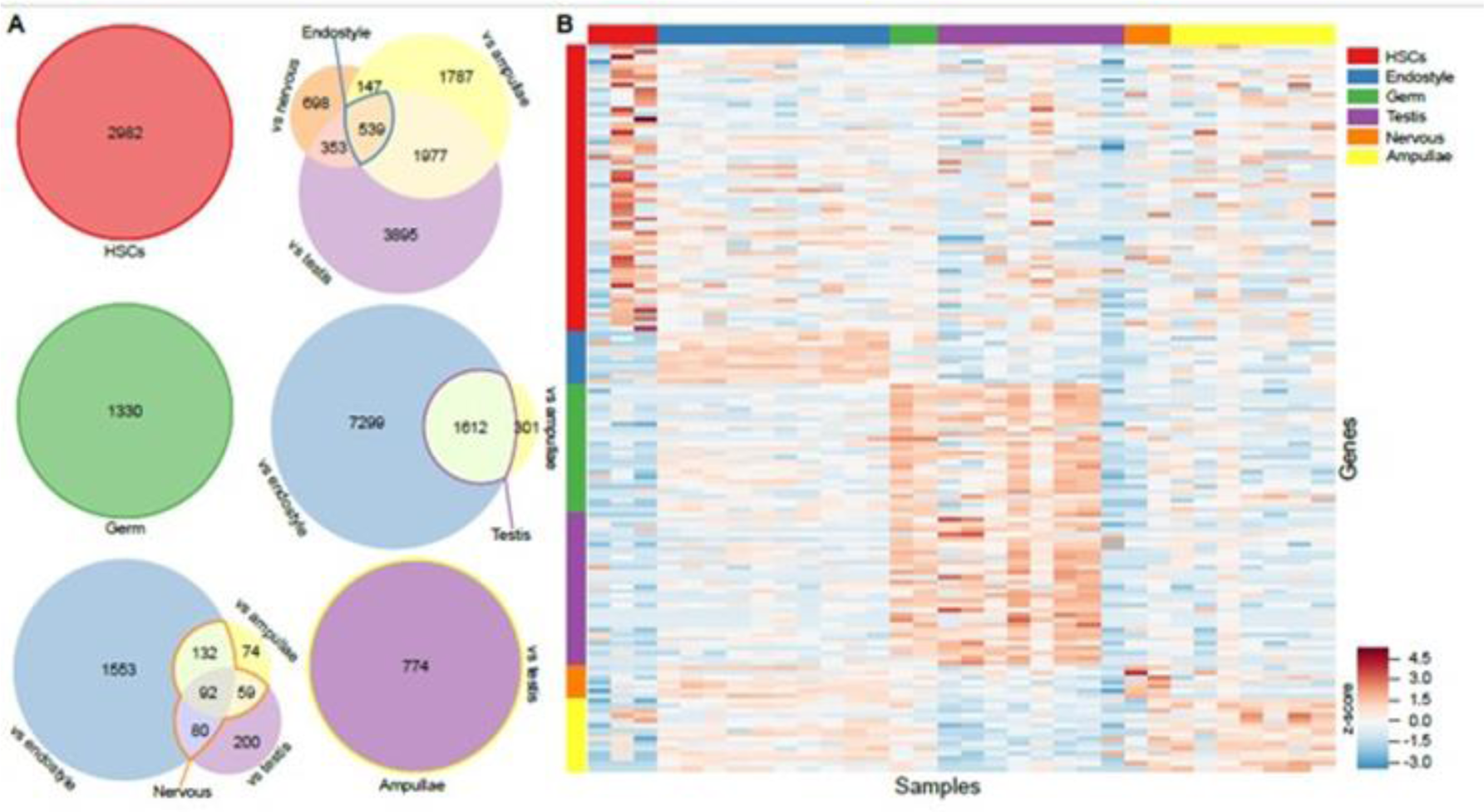
Tissue and cell-type associated genes. A) Venn diagrams showing the number of genes and which tissues were compared to select the gene set. HSC and germ cell associated genes are taken from prior work in characterizing circulating cells in *B. schlosseri* blood (Rosental et al., 2018). All other tissues were compared pairwise and various intersections of differentially up-regulated genes were selected (indicated by bold outline of regions with the label for the tissue). B) Heatmap showing expression of a subset (2%) of the enriched genes (rows) in the samples (columns) for each tissue. Values are z-scores of the log-transformed counts/million.

**Figure S3:**
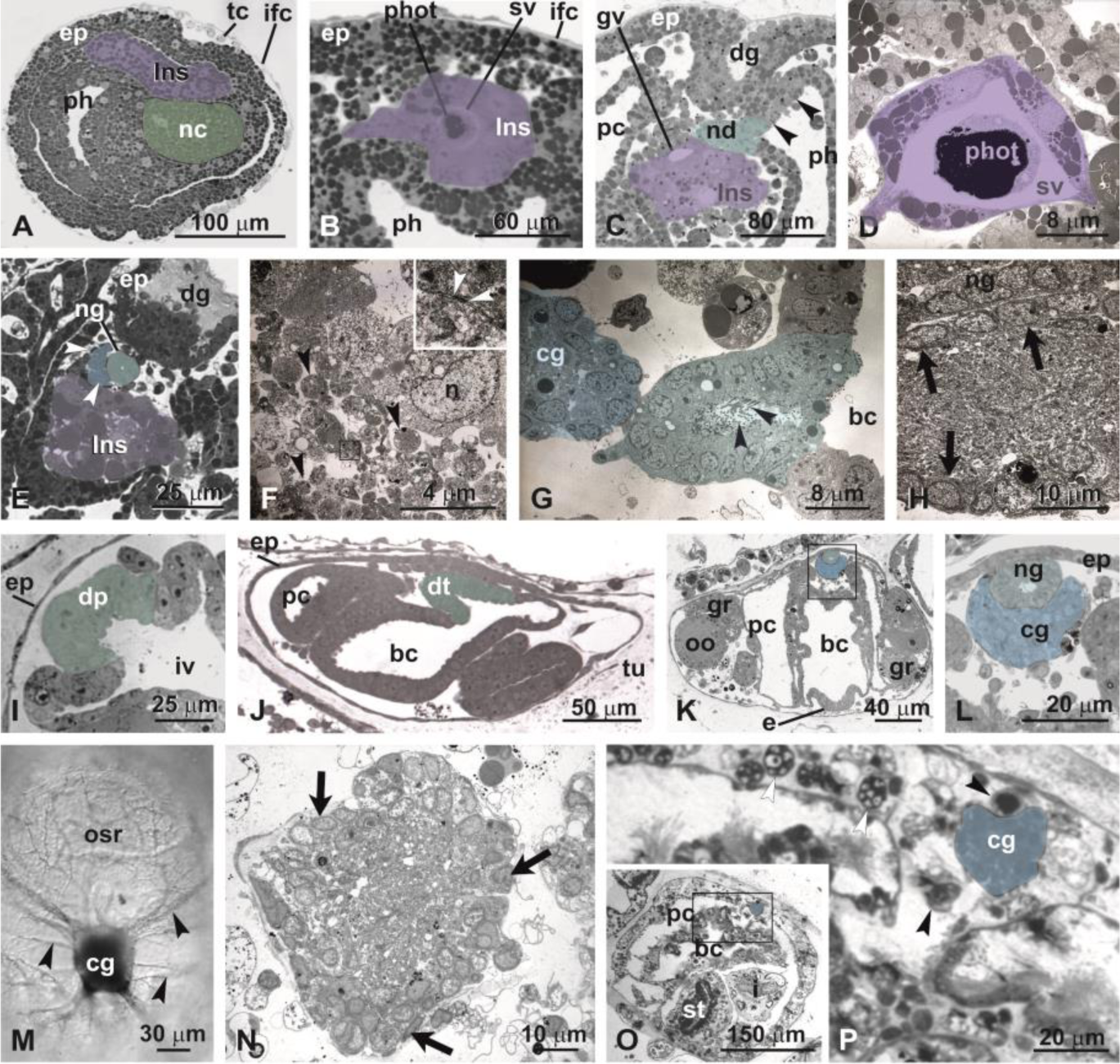
Nervous system development in embryogenesis (A-H) and blastogenesis (I-P). A) Neurula (E.3.1), the larval nervous system (lns) is dorsal to the pharynx (ph) in the trunk and to the notochord (nc) in the tail. Frontal section; details in Table S1A; toluidine blue. ep: epidermis; ifc: inner follicle cells; tc; test cells. B) Early wrap (E4), in the anterior larval nervous system (lns), the photolith (phot) is detected in the sensory vesicle (sv), sagittal medial section, stained with toluidine blue. C) Mid-late wrap (E5-E6), the neurohypophyseal duct (nd) is separated from the left ganglionic vesicle (gv) and is open into the anterior pharynx (ph). Its aperture, which is ventral to the oral siphon rudiment (arrowheads), represents the rudiment of the neural gland ciliated duct. Sagittal medial section; toluidine blue; details in Table S1A. dg: dorsal groove; pc: peribranchial chamber. D) TEM of photolith (phot) in sensory vesicle (sv) of an embryo at mid-late wrap (E5-E6). E-F) Larval settlement (E7-E8). E) transverse section showing the neural gland (ng) and the forming cerebral ganglion (arrowheads). The larval nervous system (lns) is degenerating. Toluidine blue. F) TEM of the degenerating larval nervous system, with loose neurites (black arrowheads). Some synapses are still recognizable (arrowheads in inset). n: neuron nucleus. G) TEM of the neural complex during metamorphosis (E8-E9), the neural gland aperture exhibits numerous cilia (arrowheads), the cerebral ganglion (cg) includes external cortex of neuron somata and inner medulla of packed neurites. bc: branchial chamber. H) TEM of cerebral ganglion in an oozooid. The cortex is formed of one to two layers of neuron somata (arrows), and the inner medulla has packed neurites. I) Transverse section of a secondary bud (b5), focusing on the thickening of the epithelium of the inner vesicle (iv) that form the dorsal placode (dp). Toluidine blue. J) Sagittal medial section of a secondary bud (b6). The dorsal placode has formed the dorsal tube (dt), an evagination in the form of a blind tube. Toluidine blue. tu: tunic. K-L) Primary bud (B1). The cerebral ganglion (cg) is in continuity with the neural gland (ng), actively proliferating neuroblasts. The square area in (K) is enlarged in (L). Transverse section; toluidine blue. e: endostyle; gr: gonad rudiment; oo: oocyte. M) The primary bud (B5) central nervous system (cerebral ganglion, cg) connected to its nerves (arrowheads). Whole mount; acetylcholinesterase stain; light microscopy. N) TEM of the central nervous systemof an adult individual. Arrows: neurons. O-P) Regressing adult individual (TO; see histological section in Table S1B). The zooid is contracted; the cerebral ganglion (cg) is small and surrounded by large phagocytes (black arrowheads) and cytotoxic cells (white arrowheads). The square area in (O) is enlarged in (P). Transverse section. Toluidine blue. i: intestine; st: stomach.

**Figure S4.**
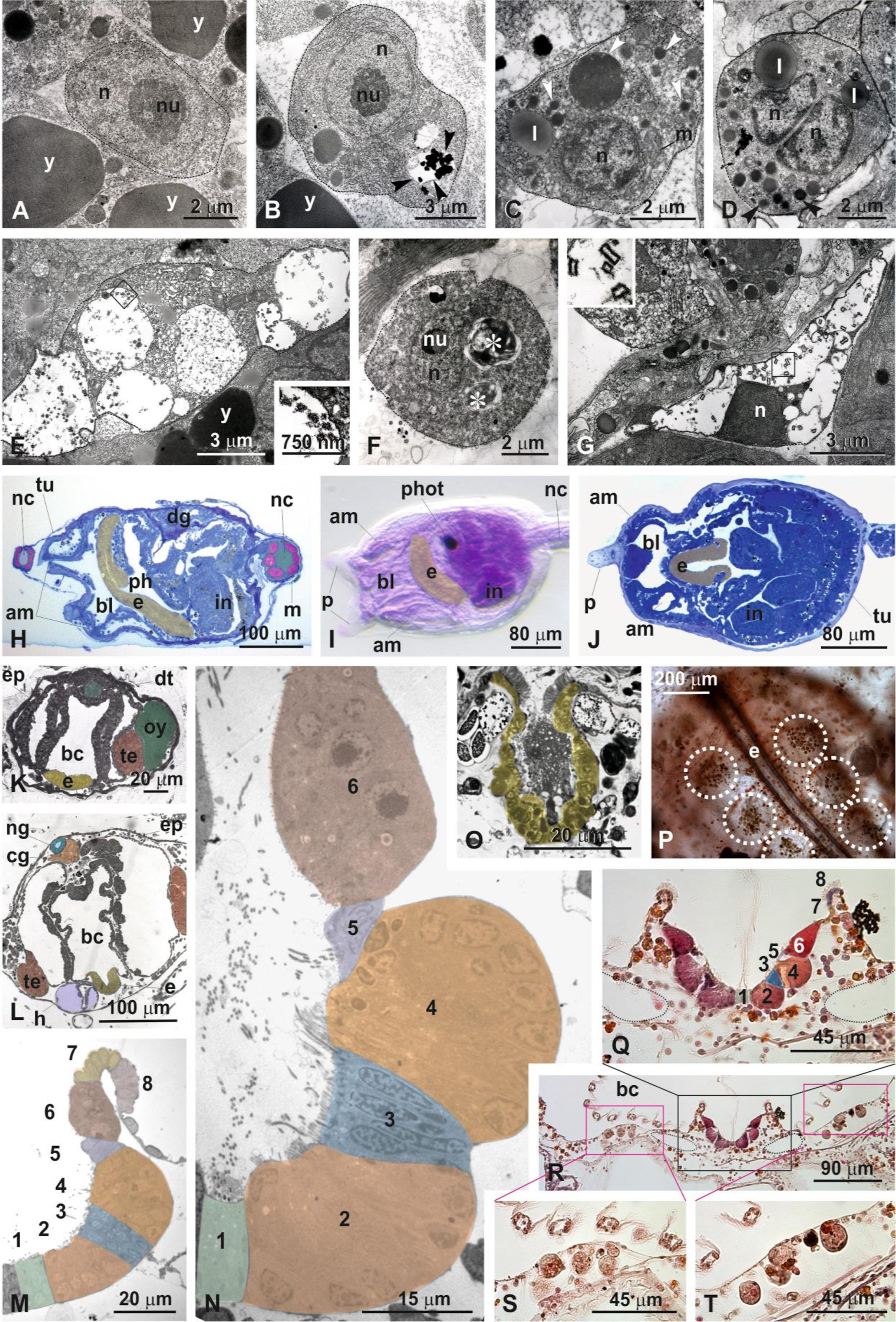
Development of the hematopoietic system and the endostyle. A-B) Early wrap (E4), TEM. A) Lymphocyte-like cell (dotted line), with high nucleus-cytoplasm ratio. n: nucleus; nu: nucleolus; y: yolk granule. B) cytotoxic morula cells, characterized by electron-dense content in vacuoles (arrowheads). C-E) Mid to late wrap (E5-E6). C) Morula cell precursor (granular amoebocyte). In the cytoplasm, there are homogeneous, membrane-bounded granules (reaching 1.5 μm in diameter) (arrowheads). D) Phagocyte (hyaline amoebocyte), with numerous, small, membrane-bounded granules (0.5 μm) with homogeneous content (arrowheads). The nucleus (n) is lobed. E) Pigment cell. Square area in E is enlarged in inset, to show the granules contained in large vacuole. l: lipid droplet. F) A settled larva (E8-E9). Phagocytes (macrophage-like cells). In the cytoplasm, there are digestive vacuoles (asterisks) with heterogeneous content. G) Oozooid (E9). Nephrocytes with granules (enlarged in inset) with hourglass geometry. H-J) Endostyle development during embryogenesis. H) Mid to late wrap embryo (E5-E6, sagittal section). The endostyle (e, yellow) appears as a groove in the anterior pharynx (ph). Note two sections of tail, with notochord (nc) flanked by three rows of musculature (pink, m) per side. Toluidine blue. I-J) Whole mount (I, seen from left side) and frontal section (J) hatched larva (E6-E7). Hemalum in I; Toluidine blue in J. am: blood ampulla; bl: blood lacuna; dg: dorsal groove; in: intestine; p: papilla; tu: tunic. K-O) Endostyle development during blastogenesis. K-L) Transverse sections of a secondary bud (b6) and a primary bud (B3). Note the endostyle (e) on the floor of the branchial chamber (bc). Organs are color-coded. Toluidine blue. ep: epidermis; h: heart; ng: neural gland; oy: ovary; te: testis. M. Transverse section of endostyle in a primary bud (B5). The eight zones (numbers 1-8) are differentiating. Colors in M, N and Q indicate the eight zones. TEM. N) Transverse sections of an adult zooid endostyle showing fully differentiated zones 1-6. TEM. O) Transverse section of an endostyle in a regressing adult zooid (stage TO). The typical zones are no longer detected. TEM. P) Endostyle (e) and cell islands (dotted circles) in a whole mount adult zooid in ventral view. Emallume. Q-T) Transverse sections of anterior endostyle and cell islands in an adult zooid. In R, black square is enlarged in Q to show the endostyle niche, red squares are enlarged in S and T to show the cell islands, dotted circles show the lumen of the blood sinuses close to endostyle. Haematoxylin– eosin.

**Figure S5:**
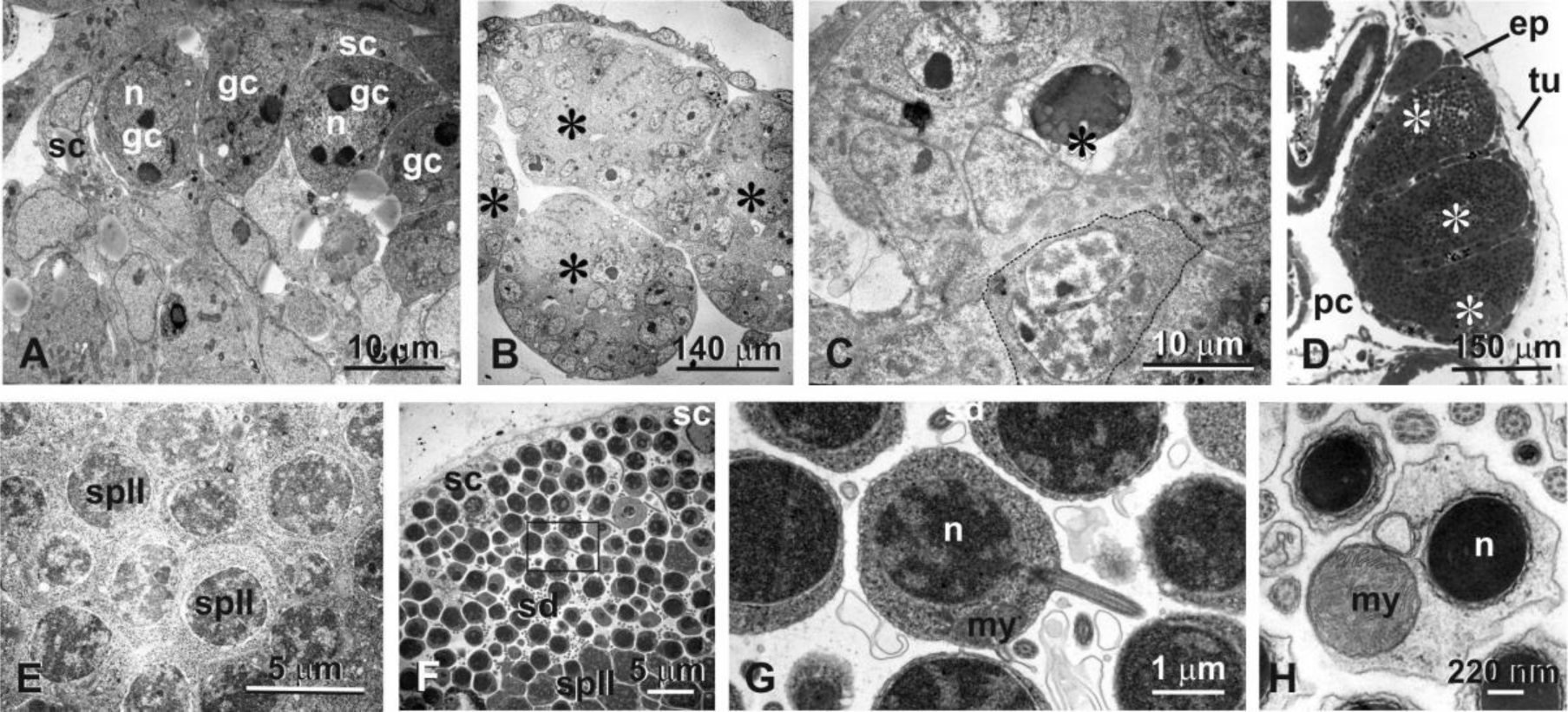
Germ cell and testis development during blastogenesis. A) Gonad rudiment in secondary bud (b3) in the form of a loose mass of candidate primordial male germ cells (gc) and candidate somatic cells (sc). Germ cells possess a nucleus (n) with one or more nucleoli and a cytoplasm with many free ribosomes, nuage, and some mitochondria. Somatic cells, which will be involved in testis wall formation, are close to germ cells. TEM. B-C) Testis in a secondary bud (b6). The testis is organized in lobules (asterisks in B). Dotted line: meiotic germ cell (primary spermatocyte). Asterisk in C: regressing germ cell. TEM. D-E) Almost mature testis in primary bud (B6). The testis is divided in lobules (asterisks in D) joining toward the sperm duct (not shown). In testis central area, numerous secondary spermatocytes (spII) are recognizable; the chromatin is condensed in the nuclei. D) Toluidine blue; ep: epidermis; pc: peribranchial chamber; tu: tunic. E) TEM. F-H) Testis in an adult zooid. F) Detail of a lobule delimited by somatic cells (sc). Spermatocyte II (spII) and spermatids (sd) are detected. Square area in F is enlarged in G to show a longitudinally cut spermatid with partially condensed chromatin in the nucleus (n), a single mitochondrion (my), and the flagellum. H) Transverse section of spermatozoa with strongly condensed chromatin. The central spermatozoon is cut at the level of the large mitochondrion (my) flanking the nucleus (n). TEM.

**Figure S6:**
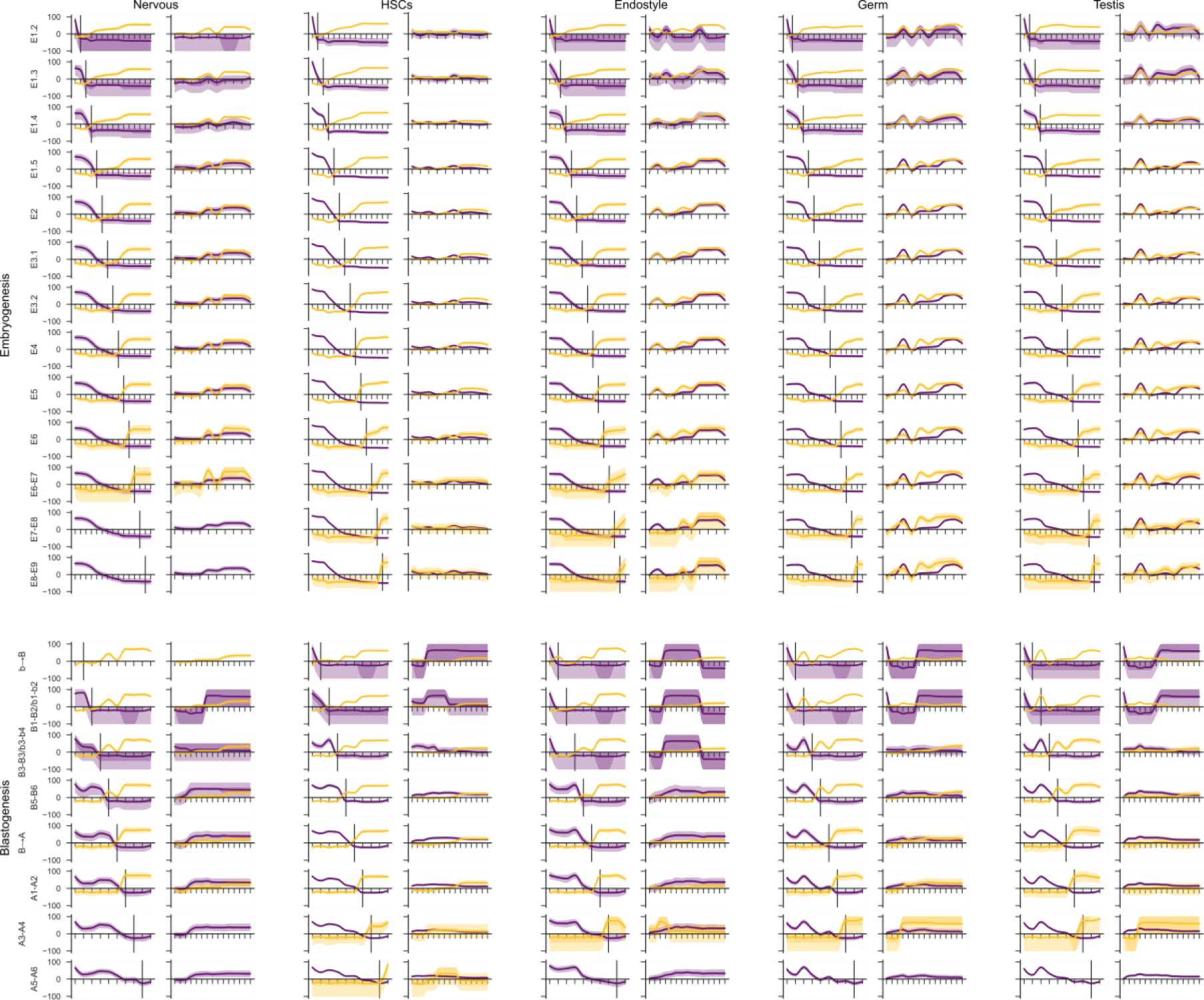
Transcriptional timelines for all tissues and cell types. Gene enrichment plots for early (purple) and late (yellow) genes in each process (top - embryogenesis; bottom - blastogenesis) associated with each tissue (each column: nervous, HSCs, endostyle, germ, testis). In each column, the left set of plots shows the enrichment of the selected genes (before/after the black vertical line) and the right set of plots shows the enrichment of these genes in the other development pathway.

**Figure S7:**
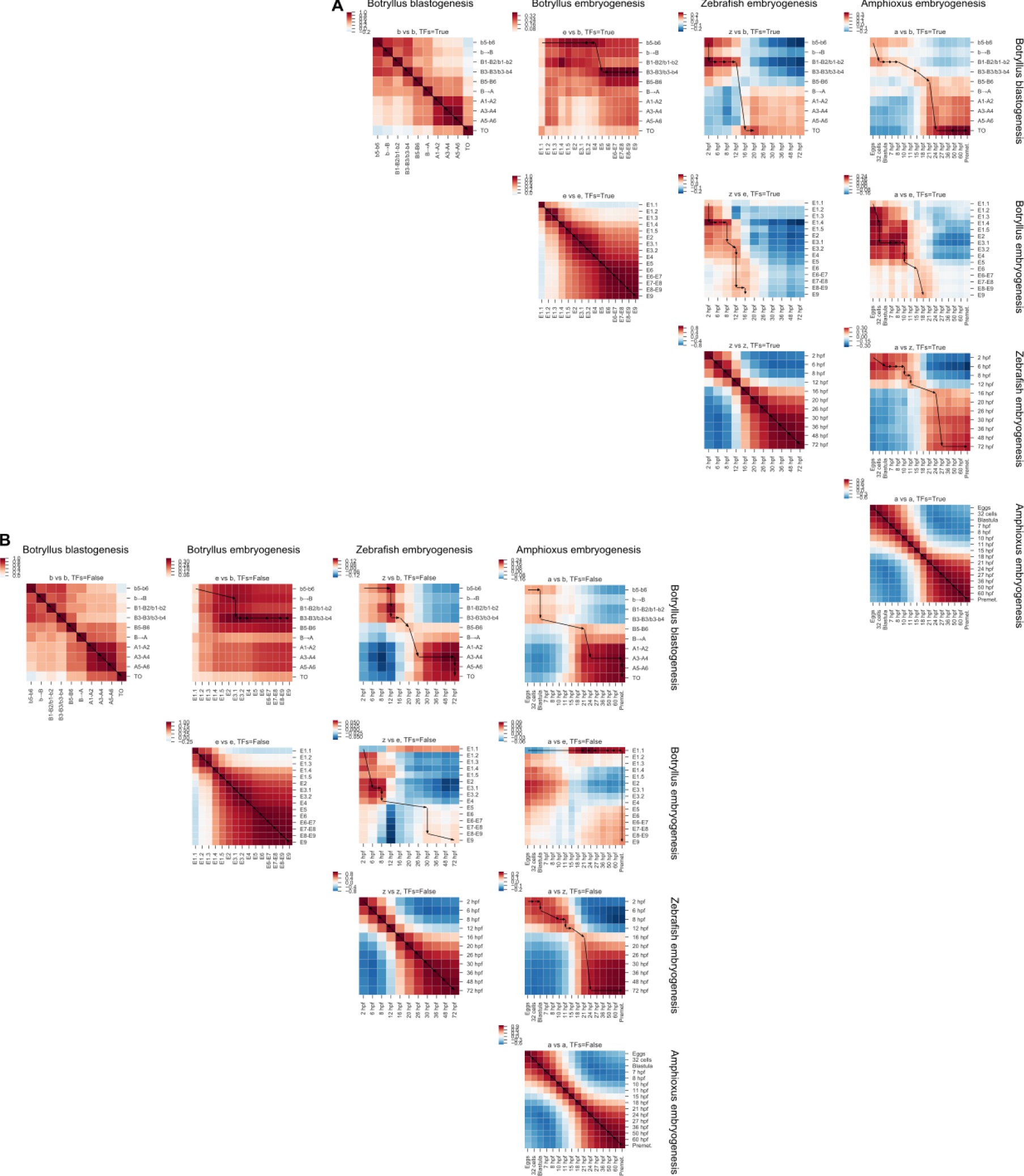
Cross species comparisons of developmental programs. Correlation plots of developmentally dynamic genes expressed at different times during developmental programs in *B. schlosseri* and two other chordates - zebrafish and amphioxus. Values range from positive (red) to zero (white) to negative (blue). Arrows and lines indicate the estimated equivalent times in the two pathways. Leading diagonals are the self-self comparison (compare with Figure 2C-D). A) Correlations restricted to putative conserved transcription factors B) Correlation over all homologous shared genes.

## STAR Methods

### Mariculture

Specimens of *Botryllus schlosseri* (family *Botryllidae*, order Stolidobranchiata) used in this study were collected from both the lagoon of Venice (IT) and Monterey Bay (CA). The colonies from Venice lagoon were used for histological analyses, 3D reconstructions and the Transmission Electron Microscopy (TEM). The colonies from Monterey Bay were used to sample specimens for confocal and two photon microscopy, RNA sequencing, and FACS analysis and to study the *ex-vivo* embryogenesis timeline.

Mariculture procedures have been described previously (Boyd et al., 1986). Briefly, wild type *Botryllus schlosseri* colonies were tied to 3×5-cm glass slides and placed 5 cm opposite another glass slide in a slide rack. The slide rack was placed into an aquarium, and within a few days the tadpoles hatched, swam to the settlement slide, and metamorphosed into the adult body plan (oozooid). Single oozooids are then transferred to individual slides and grown at 18-20°C.

Samples were collected at different blastogenesis and embryogenesis developmental stages. Staging methods were based on Sabbadin (1955) (Manni et al., 2014) staging method for blastogenesis, and the anatomical and developmental ontology of *Ciona intestinalis* (https://www.aniseed.cnrs.fr/) for early stages of embryogenesis.

For both developmental pathways a numerical staging method that refers to the day of development for embryo, secondary bud, primary bud and adult zooid was developed (Table S1; Figure S1), simplifying the comparison between *B. schlosseri*’s two developmental pathways and embryogenesis in other chordate species.

## *Ex-vivo* embryogenesis

Parental colonies were dissected under a Wild Stereomicroscope, and embryos were collected and separated into 35mm petri dishes according to stage. Embryos were kept in filtered seawater at 23C° and their development was tracked by time-lapse microscopy using a BZ-9000 Keyence microscope (Video 1).

### Sample collection for sequencing

Tissue samples were collected from *Botryllus schlosseri* colonies raised in the Hopkins Marine Station Mariculture Facility. Animals were isolated without food for 20 hours prior to dissection. Tissues were obtained by microdissection, frozen in liquid nitrogen, and held at −80 prior to library preparation.

### Library preparation

RNA was prepared from frozen samples using Zymo Research Quick RNA MIcro Prep Kit # R1050, and cleaned using Zymo Research RNA Clean and Concentrator, #R1015. Samples were analyzed on an Agilent QC 2100 Bioanalyzer to determine quality prior to library preparation.

cDNA was prepared used the Nugen Ovation RNA Sequencing System V2, #7102 and cleaned using the Qiagen QIAquick PCR purification kit, #28104, as recommended in the protocol and analyzed on the Agilent QC Bioanalyzer. If needed, samples were size-selected using Zymo Research Select-A-Size DNA Clean and Concentrator #D4080 prior to barcoding. Final library was prepared using NEB NEBNext Ultra II DNA Library Prep Kit #27645 and barcoded using NEBNext Multiplex Oligos for Illumina #E6609S. All magnetic bead purification was accomplished using BullDogBio CleanNGS RNA and DNA Spri Beads #CNGS005. Samples were then analyzed on the Agilent QC 2100 Bioanalyzer to determine the concentration of each sample prior to determine dilution prior to sequencing. On average, 12 million 2×150 bp reads (Illumina Nextseq 500) were sequenced for each library.

### Gene counts

Following sequencing, reads were processed using a Snakemake (Köster and Rahmann, 2012) pipeline: they were trimmed to remove low quality bases and primers, merged if the reads from both ends overlapped, and aligned to a database of *B. schlosseri* transcripts using bwa (mem algorithm), with likely PCR duplicates removed and then read counts determined for each transcript, resulting in 3 count tables: S4A-embryogenesis, S4B-blastogenesis, and S4C-tissue specific.

### Developmentally dynamic genes

Samples were selected and differentially expressed genes were found using edgeR (Robinson et al., 2010) from all sets of contiguous times in either pathway (Figure 1C, upper right). For each gene, all such comparisons for which statistically significant differences (FDR < 0.05) were collected and the best time signature (with zero, one or two “humps”) that explains these DE observations for each gene was found (Figure 1C, lower right). To further simplify the comparisons, these time signatures were binarized, with 1 indicating “high” expression and 0 indicating “low” or zero expression producing a gene-time expression matrix for each gene along the developmental pathway (Table S2A-B).

### Tissue enriched genes

For the nervous system, endostyle, testis and ampullae, differentially expressed genes were found on all pairwise comparisons and then gene sets picked from unions and intersections of these comparisons (see Figure S2A; Table S2D). Existing gene lists associated with HSCs and candidate germ cells (Rosental et al. 2018), were also used.

To determine time dynamics of these gene sets the proportion of these genes that are active (“1” in the binary time signature) at all times are calculated. From this value a baseline value is subtracted. The baseline calculates using a hypergeometric model the average proportion of genes that are active assuming a random selection of genes of the same size as the gene set. To determine uncertainties, the 50% and 99% confidence intervals are also calculated using the same hypergeometric model.

### Eras

The correlation of the binary developmentally dynamic gene matrix at each stage against all other stages was calculated. From this, eras of self-similar times were able to be identified. They were identified by determining the times which become less similar to the following time than they do former ones, indicating a discontinuity in the transcriptional profile and hence a transitional time (Figure 2C-D; Table S2C).

### Species comparison

To compare developmental dynamics between other chordates, public raw sequence data was downloaded from NCBI for amphioxus and zebrafish (Marlétaz et al. 2018). These were processed using a Snakemake pipeline in a similar way to the *B. schlosseri* sample. Reads were trimmed, aligned to UniVec core using bowtie2 and aligned reads removed. Cleaned reads were aligned to the appropriate reference databases using STAR (Dobin et al. 2013), duplicates removed using Picard and htseq (Anders et al. 2015) (intersection-nonempty mode, secondary and supplementary alignments ignored) against reference GTF files. From count tables developmentally dynamic genes were determined in the same manner as for *B. schlosseri*.

To compare different times in different organisms, rows (gene IDs) were grouped by gene name. If multiple gene IDs had the same name, they were collapsed into a single object with all positive values (“1”) being kept. This in general was done for all times in different developmental pathways and common gene names kept. Optionally the list of common genes was also restricted to those that shared names with a list of human transcription factors (Lambert et al. 2018). The two binary gene enrichment vectors are then compared and the correlation distance computed.

### Klee plots

In order to present ordered (by time) heterogeneous (different numbers of samples at each time point) data in a heatmap-like form, code to produce “Klee” plots (named after Swiss artist Paul Klee whose works include irregular colored rectangles). Each gene and time is given an equal spacing, with rectangles inside this region showing the expression of individual samples. The color of lines on the outside of entries is the average value over samples for a given gene and time.

### Histology

Embryos and buds were fixed for 2 hours in 1.5% glutaraldehyde in 0.2M sodium cacodylate and 1.6% NaCl buffer. After 3 washes in 0.2 M sodium cacodylate and 1.6% NaCl buffer, samples were post-fixed for 1 1/2 hours in 1% OsO4 in 0.2M cacodylate buffer at 4°C. Samples were dehydrated and then soaked in Epon and propylene solution at 37°C, 45°C, and 60°C. They were then embedded in resin, oriented and sectioned using a Leica ultramicrotome. Sections, 1μm thick, were stained with toluidine blue.

### Electron microscopy

Colonies were anesthetized with MS222 for 5–10 minutes; then, selected fragments of colonies, cut with a small blade, were fixed in 1.7% glutaraldehyde buffered with 0.2M sodium cacodylate plus 1.6% NaCl, pH 7.4. After washing in buffer and post-fixation in 1% OsO4 in 0.2 M cacodylate buffer, specimens were dehydrated and embedded in epoxy resin (Sigma-Aldrich). Semithin sections were stained with 1% toluidine blue in borax. Ultrathin sections (80 nm thick) were stained with uranyl acetate and lead citrate to provide contrast. Photomicrographs were taken with a FEI Tecnai G12 electron microscope operating at 100 kV. Images were captured with a Veleta (Olympus Soft Imaging System) digital camera.

### 3D reconstruction

An oozooid, a bud and a newly settled larva (adhesion), were embedded in resin as previously described and serially transversely cut using a Histo Jumbo Knife (Diatome). Sections, 1μm thick, were arranged in chains of about 20 sections each and stained with toluidine blue. All the sections were then photographed with Leica DMR optical microscope. Images were aligned using Adobe Photoshop CS on a Windows 7 computer. Based on the resulting stack of images, 3D models of the anatomy of all organ systems were created in Amira software (Thermofisher scientific).

### Confocal imaging

Embryos, larvae, secondary buds, primary buds and zooids were dissected under a Wild Stereomicroscope, collected and separated into 35mm petri dishes according to stage. Following several washes in sterile filtered seawater samples were fixed for 30 min at room temperature with 4% paraformaldehyde in MOPS buffer (0.1M 3-(N-Morpholino) propane sulfonic acid), adjusted to pH 7.5 and washed 2 times in 1xPBT buffer (Phosphate-buffered saline with 0.1% triton-100). Fixed samples were stained for 30 min in 1/1000 diluted cell mask orange for staining cytoplasm. After 3 washes with PBT, Alexa Phalloidin 546 was used for actin staining overnight at 4 C°. Samples were made transparent by dehydrating them with a series of solutions of 2– propanol in PBT and then with BABB (benzyl alcohol (Sigma B-1042)/ benzyl benzoate (Sigma B-6630) 1:2 ratio). In case of nucleus staining, embryos were stained with DAPI (Vector Laboratories) instead of BABB and mounted in Vectashield mounting medium. Stained samples were observed using confocal laser microscopy (Olympus fv1000) under x10 - x40 oil objective lens. 3D images were reconstructed from stack images (interval 1 to 3 μm) using Imaris software.

### Flow Cytometry

Embryos were taken at different stages and cell suspension was isolated as described (Rosental et al., 2018). Briefly, *B. schlosseri* embryos were meshed and filtered through a 40 µm mesh using a sterile 1 ml syringe pump. Cells were washed and collected in staining media: 3.3x PBS, 2% FCS and 10 mM Hepes. Cells were labeled Propidium Iodide (PI), to differentiate live vs dead cells, and with Alkaline Phosphatase (AP) Live Stain (Life Technologies A14353) 1µl, for labeling of candidate stem cell populations (Rosental et al., 2018). After gating on negative PI cells (using two dimensional plots due to natural fluorescence of *B. schlosseri* cells), the cells were analyzed on positive AP and - forward scatter (FSC) and granularity - side scatter (SSC) panel on log scale using BD ACCURI-C6. All experiments were done on pooled embryos (at least 6 individuals) for each measurement. Analysis of flow cytometry data was accomplished using FlowJo V10 (FlowJo).

### Supplementary Tables

***Tabula compositi chordati***

Table S1 - A detailed description of *B. schlosseri* embryogenesis (S1A) and blastogenesis (S1B) developmental pathways.

Table S2 - The developmentally dynamic gene lists expressed during embryogenesis (S2A), blastogenesis (S2B), eras of development (S2C) and in specific tissues and organs (S2D).

Table S3 - A detailed description of organ development through embryogenesis and blastogenesis (Table S3)

Table S4 - Transcript counts along embryogenesis (S4A), blastogenesis (S4B) and in specific tissues and organs (S4C)

Table S5 - The developmentally dynamic gene lists expressed during amphioxus (S5A) and zebrafish (S5B) embryogenesis.

### Supplementary Videos

**Supplementary video 1: *B. schlosseri* embryogenesis,** Time-lapse acquisition of *B. schlosseri* embryogenesis. In these time-lapse images we followed the development of a zygote that over the next five days (24C°), became a swimming larva. That larva features chordate characteristics such as a tail, notochord, neural tube and segmented musculature. Upon settling, the larva metamorphoses into an invertebrate individual, an oozooid (zooid originating from oocyte) which already carries both primary and secondary buds as the precursors for the next zooid (blastozooid, zooid originating from buds) generation. Taken on a Kyence BZ-X700. B-primary bud, amp-ampulla, H-heart, ds-digestive system, end-endostyle.

Supplementary **video 2: *B. schlosseri* blastogenesis,** Time-lapse acquisition of the budding cycle in *B. schlosseri*. In these time-lapse images we followed the growth and the development of a bud and the replacement of a zooid during a 6 day cycle (24C^O^). The movie focuses on an adult zooid: its primary buds, secondary buds, and the common vasculature. At the periphery of the colony, the vasculature ends in sausage-shaped protrusions called ampullae, which contract in a synchronized fashion, and act as auxiliary hearts. The zooid heart, digestive system, endostyle and the circulating blood cells can be seen clearly through the transparent body of the zooid and the colony vasculature (ventral side). The last part of the video clip shows the primary bud growing into an adult zooid, taking over the place of its parent, which is destroyed through a massive wave of apoptosis (resorbing zooid). This asexual budding cycle continues throughout the entire life of the colony (images were taken every 30 minutes during 6 days). Z-zooid B1 and B2-primary buds, sb-secondary bud,,amp-ampulla, H-heart, ds-digestive system, end-endostyle, rZ-resorbing zooid.

